# ResolVI - addressing noise and bias in spatial transcriptomics

**DOI:** 10.1101/2025.01.20.634005

**Authors:** Can Ergen, Nir Yosef

## Abstract

Technologies for estimating RNA expression at high throughput, in intact tissue slices, and with high spatial resolution (spatial transcriptomics; ST) shed new light on how cells communicate and tissues function. A fundamental step common to all ST protocols is quantification, namely segmenting the plane into regions, each approximating a cell, and then collating the molecules inside each region to estimate the cellular expression profile. Despite many advances in this area, a persisting problem is that of wrong assignment of molecules to cells, which limits most current applications to the level of a priori defined cell subsets and complicates the discovery of novel cell states. Here, we develop resolVI, a model that operates downstream of any segmentation algorithm to generate a probabilistic representation, correcting for misassignment of molecules, as well as for batch effects and other nuisance factors. We demonstrate that resolVI improves our ability to distinguish between cell states, to identify subtle expression changes in space, and to perform integrated analysis across datasets. ResolVI is available as open source software within scvi-tools.

## 1 Introduction

Spatial transcriptomics (ST) provides a powerful tool for studying the structure and function of tissues [1]. A rapidly evolving class of ST technologies operates at a resolution lower than the size of a cell, thus allowing a representation of the individual cells that comprise a tissue, along with their RNA expression profiles. These representations offer new opportunities for studying how cells operate in their tissue context and, through intercellular interactions, shape tissue function. Technologies for high-resolution ST are broadly divided in two: one set of methods rely on RNA sequencing of RNA bound to sub-micrometer spatially barcoded spots, and the other rely on imaging individually-tagged RNA molecules. To obtain a cellular resolution view of tissues from these measurements, cellular segmentation has to be performed. Namely, the tissue is divided into small regions, each representing a cell, and the molecules inside each region are counted to estimate the expression of every gene in each cell.

Early algorithms used for segmentation of ST (including legacy methods from microscopy) relied on images of the tissue under investigation that outline individual nuclei or cell membranes [2, 3]. Despite recent advances in this area, correctly defining cell boundaries remains difficult, with some of the main caveats including high cellular density, weak signal of cell staining, and irregular cell shapes [4]. As a result, the inferred segments may not correctly distinguish the number or shapes of the cells. This results in subsequent errors in expression quantification with erroneous expression estimates (e.g., suggesting co-expression of marker genes from different cell lineages).

More recent methods aimed to address this by including observed RNA molecules as an additional source of information to help guide segmentation [5, 6]. Although this coupling of segmentation and quantification has largely improved accuracy, the above problems persist (as we also demonstrate in this manuscript). In fact, in addition to difficulties in segmentation, there are caveats that directly pertain to the quantification step, with some molecules appearing within the boundary of the wrong cell. This so-called diffusion phenomenon may result from leakage of RNA molecules due to issues with tissue handling, or from overlap of cells in the third (perpendicular to the profiled plane) dimension, which is normally profiled at a low resolution and to a restricted (normally tens of microns) width.

To address these challenges, we developed resolVI, a probabilistic framework for representation of ST data while accounting for errors in segmentation and quantification. Instead of tackling the problem de novo, resolVI operates downstream of any algorithm for segmentation and quantification, taking initial estimates of gene expression in cells as its input. At its basis is the realization that diffusion of signals is an artifact that should be characterized and then controlled for. Relying on an autoencoding variational Bayes, resolVI provides artifact-corrected and probabilistic estimates of both low dimensional representation of cells as well as their gene expression profiles. Through a number of test cases for image- and sequencing-based technologies, we demonstrate that resolVI substantially improves our ability to identify cellular populations. Additionally, we demonstrate that resolVI is capable of eliminating erroneous co-expression patterns while at the same time preserving unexpected co-expression patterns reflecting nuanced cell populations. Our probabilistic model enables robust hypothesis testing of differential expression and cell colocalization. ResolVI is available as an open source project under scvi-tools [7].

## 2 Results

### Overview of resolVI

The input to resolVI is a matrix of gene expression in cells, their respective spatial position, and the respective tissue slice (in case there are multiple tissue slices). These matrices are generated from the raw ST data (RNA sequencing of spots or positions of individual molecules) by any quantification algorithm. The output of resolVI is a probabilistic low-dimensional representation of the state of each cell corrected for batch effects, along with estimates of its expression profile corrected for artifacts.

To achieve this, resolVI employs a latent variable model inferred with a variational auto-encoder (VAE). The model represents the observed counts as the sum of three components: true expression in the respective cell, expression that originates from nearby cells, and expression due to an unspecific background (constant for each tissue slice). The expression in cells (the first two components) is modeled using the negative binomial distribution [8]. For the third component (background), we assume no overdispersion and model it with a Poisson distribution (Figure 1A, B). The counts in each cell are generated from a low-dimensional latent representation that represents its cellular state. Our uncertainty about cell states is expressed using a Gaussian mixture distribution. Unlike the standard unimodal approach used in VAEs, this choice helps to better distinguish between cell types and provides users with the option to include predetermined cell-type labels during training [9, 10].

**Figure 1:**
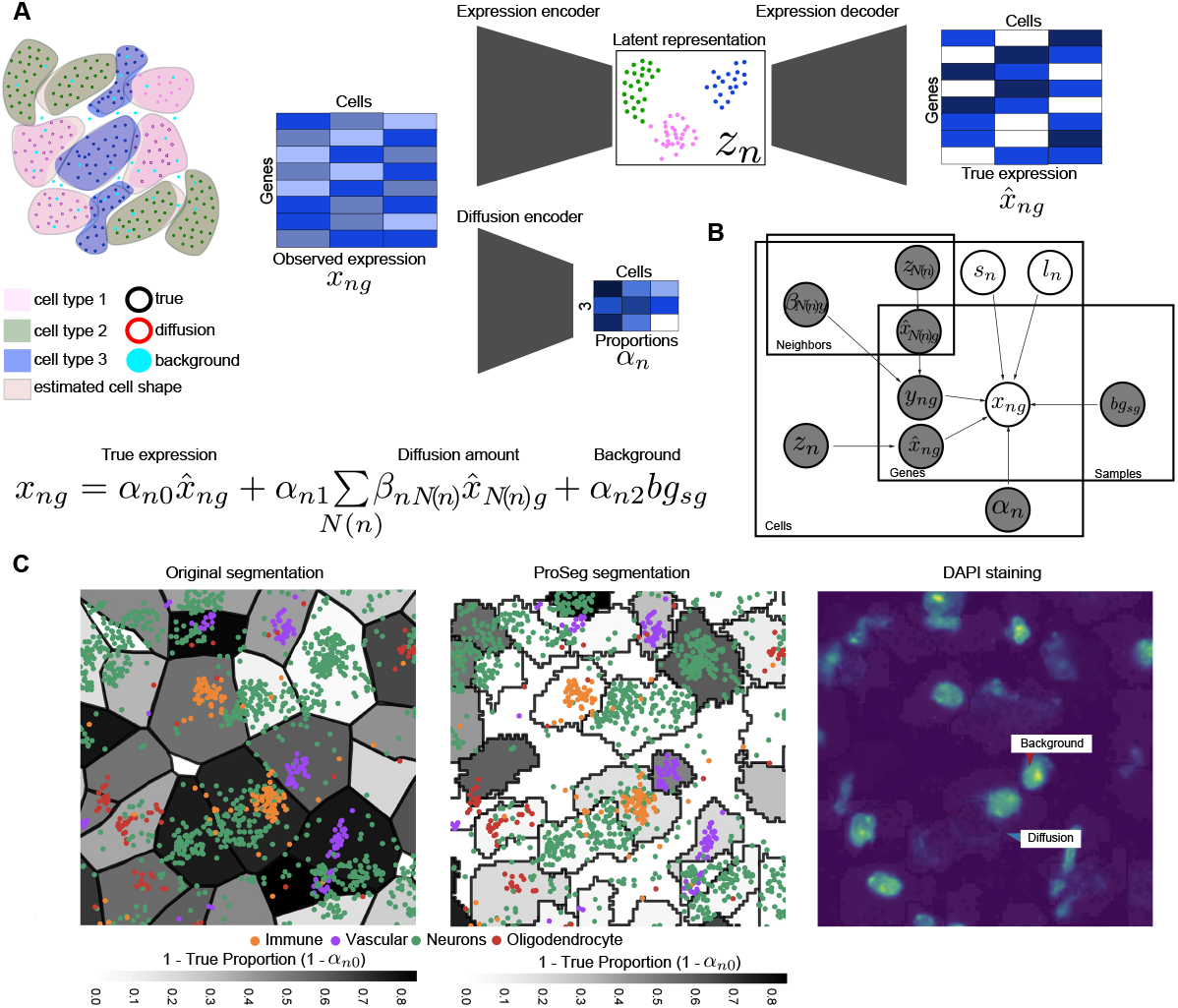
Overview of resolVI. **A** Schematic of algorithm. To assign molecules to cells in spatial transcriptomics segmentation is performed. These algorithms do not fully capture the true cell shape. This leads to misassignment of molecules. Additionally, unspecific background is present. Both phenomena contribute to the observed expression. Latent code of a cell and contribution of “diffusion” and “background” to observed expression are estimated per cell using encoder networks. The latent code is decoded to yield the true expression. The mean observed expression is reconstructed as a sum over the estimated true expression, a weighted sum over the estimated true expression of neighboring cells to reconstruct diffusion and a per sample background gene expression. We use a Poisson distribution for the background and a negative binomial distribution for diffusion and true expression (see Methods). **B** Plate model of resolVI. *z*_*n*_ and *z*_*N*(*n*)_ are the latent codes of the center cell and the neighboring cells respectively with *N* (*n*) being the neighboring cells of cell *n. β*_*nN*(*n*)_ is the per neighbor contribution and 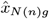 denotes the gene expression of cells *N* (*n*). *s*_*n*_ and *l*_*n*_ are the sample ID and molecule count of cell *n*. 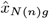, and 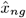 depend on the molecule count and sample ID. The arrows are not displayed to simplify the plate model. *y*_*ng*_ is the total estimated gene expression due to diffusion summed over neighboring cells *N* (*n*) and *bg*_*sg*_ is the estimated background of sample *s*_*n*_. Grey plates are latent parameters, while transparent ones are observed. In the semi-supervised model *z*_*n*_ and *z*_*N*(*n*)_ are dependent on the (partially) observed cell-type label of the respective cell-type label *c*_*n*_ and we additionally train a cell-type classifier that predicts *c*_*n*_ from *z*_*n*_. **C** Highlighted is a region of the mouse brain recorded with Xenium. Displayed are the original 10X provided (left) and ProSeg (middle) segmentation. Molecules of cell-type marker genes are colored by their respective cell-type. Black lines are segmentation boundaries and cell shapes are colored by estimated true proportion. We show the respective DAPI image on the right. We highlight one case of diffusion, where the boundary between immune cell and neuron is not clear and one case of background, where a vascular cell contains sparse neuron-related transcripts.

The model is estimated by optimizing the evidence lower bound and consists of three neural networks: an expression encoder, which estimates cell state from the observed counts, an expression decoder, which generates true expression given a cell’s state, and a diffusion encoder, which estimates the weight of each of the three components (true expression, neighbors, background) for each cell. In particular, the decoder network accepts the batch ID of each cell as input, while the encoder by default does not, helping to mitigate batch effects as previously done [8]. The intuition for why resolVI is capable of distinguishing true from erroneous counts is in its implicit expectation that the data can be described with a low-dimensional encoding [8]. Due to errors in quantification, the observed expression profiles can become more complex than the true expression profiles, and therefore minimizing the ELBO loss encourages assignment of “unrelated” molecules to the background component or to neighboring cells.

Our implementation of resolVI was designed to utilize graphics processing units (GPUs) for accelerated training. All experiments were performed on a workstation using an Nvidia 3090 GPU and 128 GB of memory. In our largest test case, which included 1.4 million cells from 52 slices [11] (Figure 5), the training time was just under six hours. For comparison, segmenting a single slice of a medium-scale experiment requires anywhere between several hours using ProSeg and several days using Baysor. Including resolVI in a workflow therefore adds a small overhead to the overall computational burden.

### ResolVI improves cell-type resolution and corrects spurious co-expression patterns in mouse brain data

As our first case study, we use a single slice of a normal mouse brain, profiled using the fluorescence-based 10X Xenium technology. This data set consists of a panel of 248 genes and was segmented by its original authors into 130k cells. The segmentation and quantification procedure was based on nuclei segmentation and subsequent dilation; this divides the tissue into convex polygons that surround each nucleus. Molecules inside each polygon are counted to estimate the respective expression levels. Since the data came without cell type labels, we applied an annotation strategy that relied on manual curation of cell type markers and label transfer (Methods, Supplementary Figure 1).

We applied resolVI on the observed expression counts using two modes. The unsupervised mode used only the gene expression data and cell locations. The supervised mode accounts for the cell type labels by linking the modes of the Mixture-of-Gaussian prior with the different cell types and adding a cell-type classifier that takes the latent space coordinates as features to predict the cell-type (Methods). A visualization of the latent representations inferred by the unsupervised resolVI model demonstrates that it captured the distinction between cell types well, especially compared to the analysis of principal components (PCA) of the observed expression counts (Figure 2A). We quantified this using the scib-metric bio-conservation measures [12] (Figure 2B). Notably, scib-metrics also offers a way to evaluate the extent of batch effect correction. Since this is not applicable in this case (as the data consist of one slice), we divided the cells into low and high estimated rates of erroneous expression per resolVI. To this end, we define a cell as high-rate when the proportion of molecules assigned to the diffusion or background components (i.e., *α*_1_ + *α*_2_ in Supplementary Figure 1) is larger than 20%. We would expect a good integration method to pull together cells of the same type, despite differences in the rate of erroneous molecule assignment (as a proxy for data quality). We found that the addition of supervision improves the ability of resolVI to integrate low- and high-quality segmentations. Furthermore, resolVI with and without supervision compared favorably to scANVI (which also uses supervision), scVI, and PCA in both bioconservation and batch correction (Figure 2B and Supplementary Figure 2). For the remainder of this data set, we analyze the model without cell-type supervision, as this is a more common scenario.

**Figure 2:**
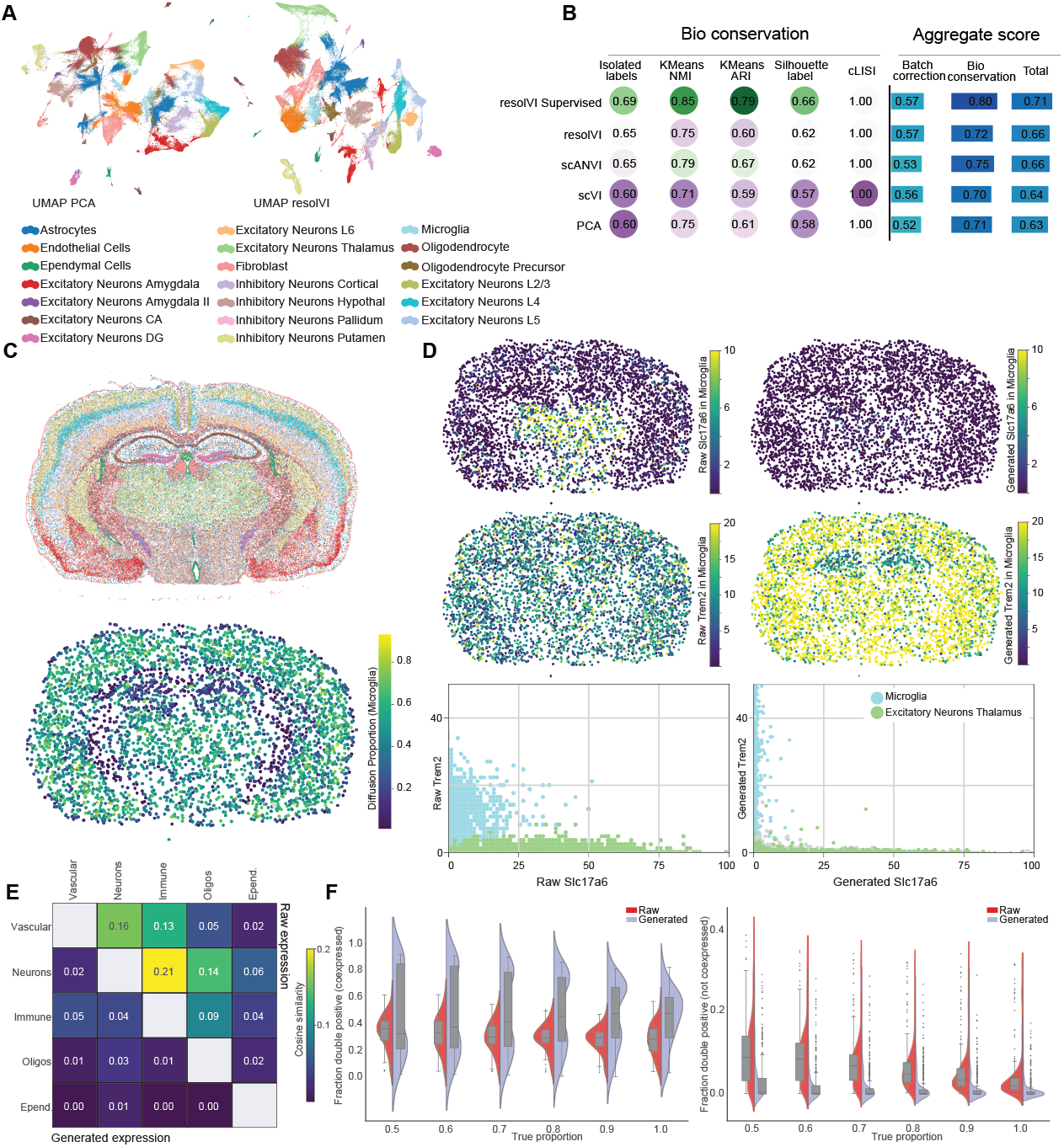
ResolVI corrects expression biases in mouse brain. **A** UMAP embedding of PCA computed on count normalized expression data (right) and resolVI latent space) using 10X provided original cell segmentation. Displayed are our manual cell-type labels. **B** To quantify these findings, we computed scib-metrics on these embeddings. As batch labels, we use all cells with an estimated true proportion above 0.8 (empirically determined) to benchmark whether low and high quality cells are integrated. We show a subset here and only display the Batch Correction aggregation score (full metrics in Supplementary Figure 2). **C** Spatial positions of cells are colored by cell-type labels (top) as well as all cells labeled as microglia colored by their diffusion proportion (bottom). **D** Only microglia cells are displayed in space. On the left side we show the raw expression, while on the right side we show the resolVI generated expression. *Slc17a6* is a marker of neurons, while *Trem2* is a marker of microglia. We generated expression with resolVI with the same counts per cell as the raw cells and display here the count-normalized data. The bottom row highlights a scatter plot of both genes. In these scatter plots, we display excitatory neurons of the thalamus, that express *Slc17a6* and microglia, that express *Trem2*. Spots are colored by cell-type label. **E** We calculated mutually exclusive expressed genes across cell-types using a single-cell reference dataset (Supplementary Figure 3). We display here the cosine similarity of the sum over all marker genes of the respective cell-type. Above the diagonal we display uncorrected data, while below the diagonal, we display resolVI generated expression. Lower values display less cross-contamination. **F** We use a Poisson mixture model on count-normalized data to determine marker-positive cells. (Left) Pairs of genes that are co-expressed in the single-cell sequencing dataset are displayed and (right) pairs of genes that are expressed in different cell-types are displayed. We split the cells into equal spaced bins of estimated true proportion (1.0 means all cells with estimated true proportion between 0.9 and 1.0). In both plots, we display on the y-axis the amount of co-expression (double-positive divided by sum over single-positive cells) for each pair of genes.

ResolVI estimates for each cell a diffusion rate, namely the fraction of counts (in all genes) that originate from neighboring cells (*α*_1_ in Figure 1). Taking into account their spatial distribution, we found high diffusion rates in regions with high cell density, particularly the cortex and the thalamus, probably because cell density makes these regions harder to segment (Figure 2C). To study predicted gene expression, we focus on microglia as they form several clusters before correction with resolVI (Figure 2A). These cells express *Trem2*, supporting their labeling as microglia (Figure 2D). However, in the hypothalamus, a region with high predicted diffusion, these cells are also positive for *Slc17a6*,a marker of excitatory neurons. In contrast, after running resolVI, false positive expression of *Slc17a6* in microglia is drastically reduced, while these cells are still predicted to express *Trem2*. This highlights that resolVI can reduce spurious expression patterns (likely caused by errors in delineating cell boundaries during segmentation), while preserving correct expression patterns.

To more broadly quantify this, we used a single cell RNA-seq data set from the mouse brain [13] to identify genes that are specific to different cell types and therefore are not expected to be expressed by the same cell (Supplementary Figure 3). We evaluated the extent of erroneous double positives for each pair of cell types using the cosine similarity of the normalized sum over the respective cell-type marker genes (Methods). We find that the corrected data generated by resolVI includes substantially fewer erroneous double positives compared to the raw counts (Figure 2E). We confirmed these findings using gene pair-level statistics and find that cells detected by resolVI as high-quality cells show a similar co-expression pattern (Supplementary Figures 4 and 5). Importantly, resolVI does not force the data to comply with preconceived notions of cell types and their markers. For example, while many erroneous coexpression patterns are removed, resolVI still predicts co-expression of endothelial markers and *Lyz2*, which is commonly a marker of myeloid cells. Endothelial cells can indeed express *Lyz2* ([14]), which was not evident in the respective single-cell RNA-seq dataset. This highlights that resolVI can recognize and preserve unexpected co-expression patterns even if those that are missed by single-cell RNA sequencing.

As another validation, we explored the relationship between the extent of spurious signal inferred for each cell (i.e., *α*1+ *α*2), and the level of co-expression of markers of the same cell type (true positive pairs; Figure 2F, left) or different cell types (false positive pairs; Figure 2F, right). Overall, considering pairs of genes that are expected to be co-expressed, we find an equal rate of double positives between raw and corrected counts and no correlation with the estimated rate of spurious signal. In contrast, for pairs that should not be co-expressed, we find a clear decrease in co-expression after applying resolVI. Finally, we find that resolVI’s estimates of the extent of spurious signal (which is done without knowledge of cell-type markers) clearly increases with the amount of false double positive (which is based on knowledge of cell-type markers; red violins on Figure 2E, right).

### Evaluating resolVI under different segmentation scenarios

The choice of segmentation algorithm can markedly influence the extent of misassignment of molecules to cells (Figure 1C). Here, we investigate the effect of the choice of segmentation algorithm on the quality of the downstream analysis with resolVI.

Starting with the brain section of Figure 2, we added three segmentations, using Baysor [5] and ProSeg [6], which both use gene expression information for better delineation of boundaries between cells, as well as a nuclei segmentation using the 10X segmentation pipeline without any additional dilation. We transfer the cell-type labels from Figure 2 to annotate the cells generated by each segmentation (Methods) and then quantify the extent of misassignment using lists of gene pairs that are expected or not expected to be expressed by the same cell-type (Supplementary Figure 6). We find that the choice of segmentation algorithm has a substantial effect, with expression-aware algorithms generating lower rates of erroneous double positives. However, this apparent misassignment still remains present. Applying resolVI downstream to each segmentation, we find that it clearly reduces the remaining amount of erroneous double positives in both cases (Supplementary Figure 6 and 7).

To further establish this, we added a second, more complex use case focusing on cancer biopsies. We use a dataset of two liver cancer biopsies [15] from two patients that were profiled using the Vizgen MERSCOPE platform. This data set contains roughly 1.5 million cells and a panel of 550 genes. We consider six segmentation algorithms as alternatives to processing this data. Four of the methods do not use gene expression data, including the original segmentation by Vizgen MERSCOPE and three settings of Cellpose implemented within the Vizgen postprocessing tool, using cell segmentation in a single or three z-layers as well as nuclei segmentation in three z-layers (Methods). As the additional two methods, we used the expression-aware Baysor and ProSeg. Since cell type labels were not provided along with this dataset, we manually annotated ProSeg segmented cells using marker genes (Supplementary Figure 8) and then transferred these labels to all other segmentations (Methods; Suppl Figures 9).

Regardless of the segmentation algorithm, we find that the integration between the two tissue sections and the distinction between cell types were clearly improved when comparing the latent space of resolVI with the embeddings calculated by PCA or Harmony [16] (using scib-metrics). This is especially prominent when considering the distinction between closely related cell types, such as B and T lymphocytes (Figure 3A, Supplementary Figure 9 and 10). In addition, we find that both expression-aware segmentations improve the distinction between cell types, both before and after applying resolVI, indicating that combining ample segmentation with resolVI provides a good strategy.

**Figure 3:**
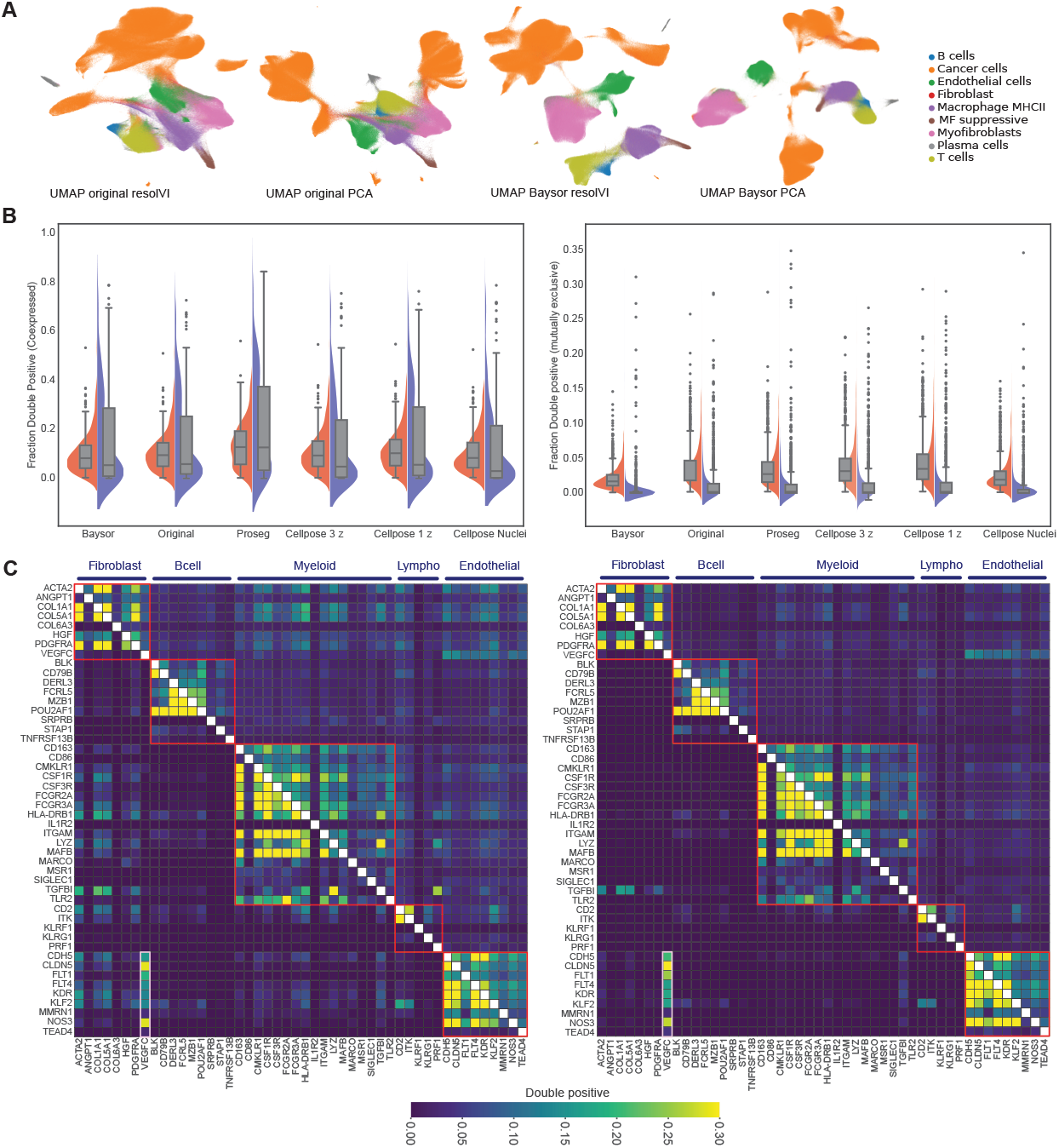
Performance of resolVI is dependent on the initial cell segmentation. **A** UMAP embedding of cells after original segmentation (left two plots) comparing resolVI (first plot) and PCA on count-normalized and square root transformed data (second plot) as well as Baysor segmentation (right two plots). All plots are colored by cell-type information. Baysor segmentation improves clustering of annotated cell-types and especially B cells cluster more distinctly in resolVI latent space compared to PCA. For quantification, we performed scib-metrics (Supplementary Figure 10). **B** Comparing the fraction of double positive cells in co-expressed genes (left) and mutually exclusive expressed genes (right). We compare the fraction before and after resolVI correction using six different segmentation algorithms. **C** Gene-gene co-expression plot for cell-type marker genes. Red blocks highlight genes that are expressed in the cell-type and potentially co-expressed while gene pairs outside of these blocks are not expected to be co-expressed. On the right side, we show the results for the original segmentation and on the left side using Baysor segmentation. ResolVI generated counts are displayed below the diagonal while above the diagonal uncorrected counts are displayed. We highlight *VEGFC* co-expression with markers of endothelial cells using a white box.

Next, we estimated the rates of true and erroneous double positives in each quantification of this data set using a single-nucleus sequencing data set from a healthy liver as reference [17]. Before adding resolVI, we found that the different segmentations yielded similar fractions of true double-positive rates, with the expression-aware Baysor showing the lowest rate of erroneous double positives. The application of resolVI reduced the rate of double positives in all cases to a comparable level (Figure 3B).

To further dissect this, we considered double positive rates for individual gene pairs (both true and erroneous) in the Baysor and original segmentations (Figure 3C). In the original segmentation, we find that resolVI led to a marked reduction in erroneous double positives. However, erroneous co-expression still remains, e.g., with fibroblast markers co-expressed with markers of other lineages, especially myeloid and endothelial. When comparing the observed expression with the original segmentation with the estimates after Baysor segmentation, we find that these co-expression patterns are largely reduced, and become yet less abundant after application of resolVI. This highlights that the accuracy of the correction by resolVI depends to a certain extent on the quality of the initial segmentation. Importantly, not all co-expression patterns that are unexpected according to the single-nuclei reference are eliminated by resolVI. For example, resolVI estimates that *VEGFC* (which appears as a fibroblast marker in the single nuclei data) is co-expressed with endothelial markers. *VEGFC* is a growth factor that acts autocrinely on lymphatic endothelial cells specifically under hypoxic conditions [18]. Hypoxia is a common feature in tumors, which explains why in addition to fibroblasts *VEGFC* is additionally expressed in endothelial cells in cancer.

ProSeg also estimates the diffusion of molecules and provides a probabilistic estimate of cellular expression. We compared these estimates with the counts generated by resolVI (Supplementary Figure 11). After ProSeg segmentation, we find a higher amount of spuriously co-expressed genes compared to the Baysor segmentation. ResolVI reduces the amount of double positive expression of mutually exclusive gene pairs, while the probabilistic estimates in ProSeg show hardly any improvement over the segmented counts. We conclude here that using Baysor as the initial segmentation step performs best for downstream analysis and will, for the remainder of the manuscript, provide results for Baysor segmentation.

### ResolVI facilitates the analysis of large sequencing-based datasets

Sequencing-based spatial transcriptomics is likely to show similar problems with spurious double positives. Due to the typically large number of genes captured in these assays, expression-aware segmentation algorithms cannot be readily applied (as they require excessive amounts of memory) and segmentation is performed with simpler methods. As our third test case, we therefore evaluate resolVI using a sequencing-based dataset of a coronal mouse brain profiled with Stereo-Seq [19] (Supplementary Figure 12). This data set includes over 24,000 detected genes and over 50,000 cells. Unlike most expression-aware segmentation methods, resolVI could be applied efficiently (runtime less than 20 minutes and memory usage below 30 GB) with all genes as input. We found a clear decrease in amount of erroneous marker gene co-expression after application of resolVI (Supplementary Figure 12b). For example, in the raw expression data, 14.8% of cells that are positive for the neuron cell-type marker *Slc17a7* were also positive for the oligodendrocyte marker *Mbp*. In the resolVI generated counts, this pattern was largely eliminated (<0.01% of Slc17a7-positive cells; (Suppl Figure 12c). Furthermore, the cell embedding and corrected counts inferred by resolVI substantially improved our ability to stratify the cells into cell-types. Microglia are difficult to distinguish in the raw data, while they are clearly distinguished in the latent space of resolVI. This is evident in the expression of marker genes such as *Trem2, Csf1r, C1qa*, and *Ctss* (Supplementary Figure 12d). Finally, the generated counts of resolVI help to distinguish inhibitory (*Gad1* and *Gad2*) from excitatory neurons (*Slc17a7*).

### ResolVI reveals functionally relevant spatial domains in human liver cancer

To evaluate resolVI in an actual analysis scenario, we applied it to a comparative study between healthy and cancerous liver samples, profiled with Nanostring CosMx [20] that included 1000 genes and 1.2 million cells. Although these data were segmented by the authors using an expression-unaware approach, we chose to reprocess them with Baysor since its combination with resolVI provided an effective strategy in our previous evaluations. We labeled the newly segmented cells by transferring the annotation provided with the original dataset (assigned with InSituType [21]), using resolVI as in previous sections (Supplementary Figure 13A) and used a cell-type supervised resolVI model for further analysis.

We first focus on the healthy liver. Hepatocytes generally exhibit a strong zonation pattern along the liver lobule, with *SAA1* marking the periportal regions, while *GLUL* marks the pericentral regions. The zonation of hepatocytes based on the difference between the expression levels of these two markers is clearly visible both in the resolVI-corrected and uncorrected data (Baysor only) (Figure 4A). In the periportal regions, we further expect to see cholangiocytes expressing *SOX4, KRT13* and *EPCAM* - marker genes that are generally expressed at lower levels than *GLUL* and *SAA1*. In the uncorrected data, we find that these lowly expressed markers are detected also in pericentral regions, while after running resolVI they have a clear periportal pattern (Figure 4B and Supplementary Figure 13B). To corroborate this, we find substantial expression of these three genes in various cell types before correction, while their expression is specific to cholangiocytes after running resolVI (Supplementary Figure 13C). This correction demonstrates the merit of accounting for background expression (*α*_2_ component in Figure 1), as this wrong signal is not derived from true expression of spatially neighboring cells since these markers are not expressed in any cell-type in the peri-central regions.

**Figure 4:**
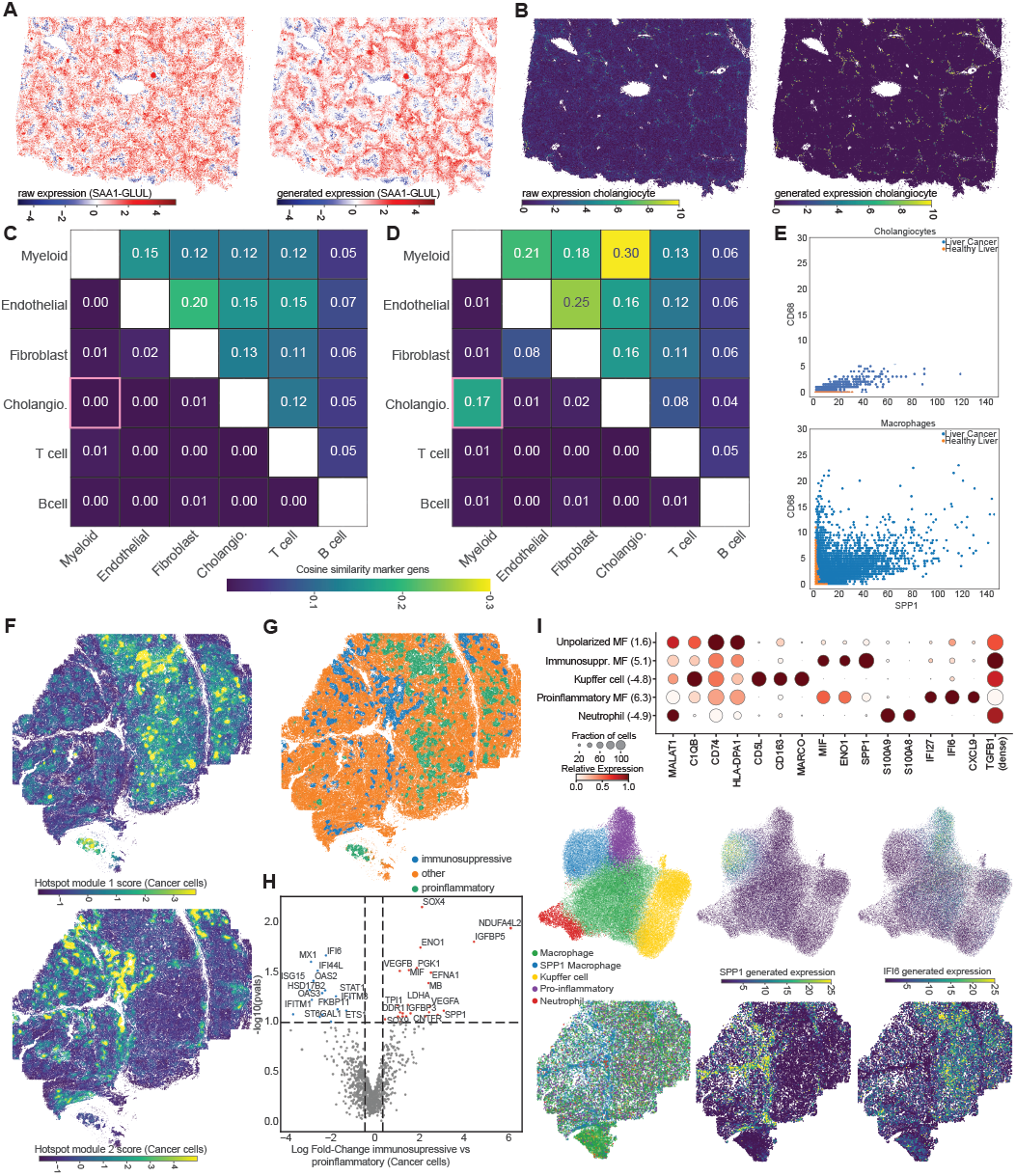
ResolVI reveals distinct niches in human liver cancer. **A** Peri-portal to peri-central gradient in hepatocytes highlighted using the difference in expression between *SAA1* and *GLUL* using raw expression (left) and resolVI generated expression (right). **B** Expression of marker genes for cholangiocytes (sum over *SOX4, KRT13* and *EPCAM*) displayed in healthy liver using original expression values (left) and resolVI generated expression (right). For both panels gene expression after count normalization is displayed. Cosine similarity computed between the sum over mutually exclusive expressed marker genes of various cell-types. Above the diagonal we highlight original expression, while below the diagonal we display resolVI generated expression values. We show this metric seperately for the healthy liver (**C**) and liver cancer (**D**). In liver cancer, we see an increase in marker overlap between cholangiocyte and myeloid markers. **E** Scatter plot of cholangiocytes (top) and macrophages (bottom). Displayed is *SPP1* on the x-axis and *CD68* on the y-axis. In orange we highlight the expression in healthy liver and in blue in liver cancer. Displayed expression values are resolVI generated expression values. Macrophages in liver cancer co-express both molecules, while in healthy liver these markers are expressed mutually exclusively. **F** Two hotspot modules were computed on cancer cells in space. Both module scores are displayed in space for the liver cancer sample. Displayed are only cancer cells. **G** A threshold was defined to define module 1 positive cells (pro-inflammatory) and module 2 positive cells (immuno-suppressive). **H** Differentially expressed genes are shown computed using resolVI DE functionality between immuno-suppressive and pro-inflammatory cancer cells. Top genes are highlighted. **I** Marker genes of macrophage sub-cell types. In brackets, we highlight the log2-fold enrichment in liver cancer vs. healthy liver tissue. Marker genes were manually selected. Displayed is the expression after count normalization and display is column normalized. *TGFBI* expression is displayed after adding a small pseudocount. Therefore the spot size displays that all cells express these genes. This helps here with displaying the difference in mean expression as the overall expression is low. **J** Macrophage sub cell-types and expression is displayed in UMAP (top row) and in space for the liver cancer sample (bottom row). On the right side the cell-type is displayed and especially immuno-suppressive macrophages are enriched in the immuno-supressive region. *SPP1* expression is confined to this region, while *IFI6* is high in the pro-inflammatory region. Displayed is the resolVI generated gene expression after count normalization.

Using a single nucleus sequencing reference from a healthy liver (Supplementary Figure 14) [17], we find a substantial decrease in the co-expression of unexpected gene pairs (e.g., those of myeloid cells and cholangiocytes) after correcting the healthy liver sample with resolVI (Figure 4C and Supplementary Figure 15). In contrast, when considering the cancerous sample, we find that co-expression between myeloid and cholangiocyte markers persists even after correction with resolVI (Figure 4D). Looking at individual gene pairs, this remaining signal is due to co-expression of *SPP1* and myeloid marker genes, such as *CD68* (Supplementary Figure 16). Further inspection of the corrected data (Figure 4E) highlights a unique property of the cancerous liver, in which tumor-associated macrophages express *SPP1*, which is a critical mediator of their interaction with tumor cells [22].

Given evidence for improved detection of co-expression patterns after applying resolVI, we next studied spatial heterogeneity in liver cancer cells. We first applied the autocorrelation-based hotspot algorithm [23] using spatial coordinates (to evaluate cell-cell proximity) along with resolVI-corrected counts to detect modules of spatially co-expressed genes. This analysis revealed two modules that are expressed by cancer cells in different locations of the tumor (Figure 4F and Supplementary Figure 17). To comprehensively study the differences between tumor cells in these locations, we adapt the lvm-DE algorithm (differential expression with latent variable models [10]) to the resolVI generative model (Figure 4H). We find up-regulation of anti-inflammatory genes such as *MIF, ENO1, SPP1* and hypoxia markers such as *SOX4, VEGFA* and *VEGFB* in the region of module 2. This combination highlights a hypoxic region in which cancer cells exhibit immunosuppressive functions. In the other region, Interferon-regulated genes such as *MX1, IFI6, STAT1* and *OAS2* are up-regulated, indicating an opposite trend in pro-inflammatory function.

To study the effect of these niches on the cancer microenvironment, we next focus on macrophages in healthy and cancerous livers. Sub-clustering of these cells reveals a distinction between myeloid cells in the healthy liver that express Kupffer cell markers and populations in liver cancer that express markers with immuno-supressive and pro-inflammatory function (Figure 4I and Supplementary Figure 18). The pro-inflammatory subset expresses *IFI27, IFI6* and *CXCL9*, while the anti-inflammatory population expresses *SPP1, ENO1* and *MIF*. The anti-inflammatory macrophages also express the highest amount of *TGFB1*, which is a key cytokine for immunosuppression in cancer [24]. The two subsets of macrophages are present in different regions of the tumor. The suppressive population, as well as the neutrophil population (expressing *S100A8* and *S100A9*, Figure 4I), colocalize with the immunosuppressive tumor subset expressing *SPP1*, while the pro-inflammatory macrophages colocalize with the pro-inflammatory cancer subset (Figure 4 J and Supplement Figure 19). We extended this analysis to T cells and were able to identify T cell states such as TREG and *CXCL13*-high CD8 cells enriched in the pro-inflammatory region, while the immuno-suppressive region is deprived of T cell infiltration (Supplementary Figure 20).

In summary, resolVI helped uncover spatial compartmentalization in a liver cancer sample, with distinct tumor states, immune microenvironments, and infiltration patterns (neutrophils in the immunosuppressive region and T cells in the pro-inflammatory region).

### ResolVI uncovers incomplete regeneration after DSS colitis

To benchmark resolVI in large studies, we use a recent study on Dextran Sulfate Sodium (DSS)-induced colitis, a mouse model of inflammatory bowel disease. The authors studied mice at different time points after acute exposure to DSS using MERFISH with a panel of 1348582 × 990 990 genes and a total of 1.4 cells in 52 samples. The day three samples point to initial signs of inflammation, while on day nine the samples show the highest amount of tissue damage. On day 21, tissue recovery is evident, while day 35 is described in the original manuscript as an almost fully recovered colon.

The authors have annotated the portion day zero - day 21 of this data set using three levels of granularity, with particularly high granularity of epithelial and stromal populations (Supplementary Figure 21). To scrutinize this annotation, we first trained an unsupervised resolVI model on this subset. We found that cells with less than 20 true counts as inferred by resolVI do not express most marker genes and are located at unexpected locations (Supplementary Figure 22). We therefore exclude these cells from further analysis. We additionally removed cell-type labels of cells that had a high mixture of different cell-types in their neighborhood in the unsupervised resolVI embedding (Methods).

Using the remaining high-confidence labeled cells of each cell type and providing no label for the other cells, we then trained a semi-supervised resolVI model (i.e., with label information available only for some cells). We used this model to relabel all cells using the cell-type classifier of resolVI. We confirmed that this new set of labels shows improved matching with marker genes (Supplementary Figure 23). We implemented a network surgery procedure for resolVI, similar to scArches [25] and used this procedure to integrate the day 35 samples into the reference model (without affecting the day zero - day 21 samples). Cell-type label transfer was performed using resolVI Overall, we find a good integration of the samples within each time point (especially using the supervised model; Figure 5A and B and Supplementary Figure 21), while the different time points show distinct distributions in the latent space due to the ongoing colitis (Figure 5C).

**Figure 5:**
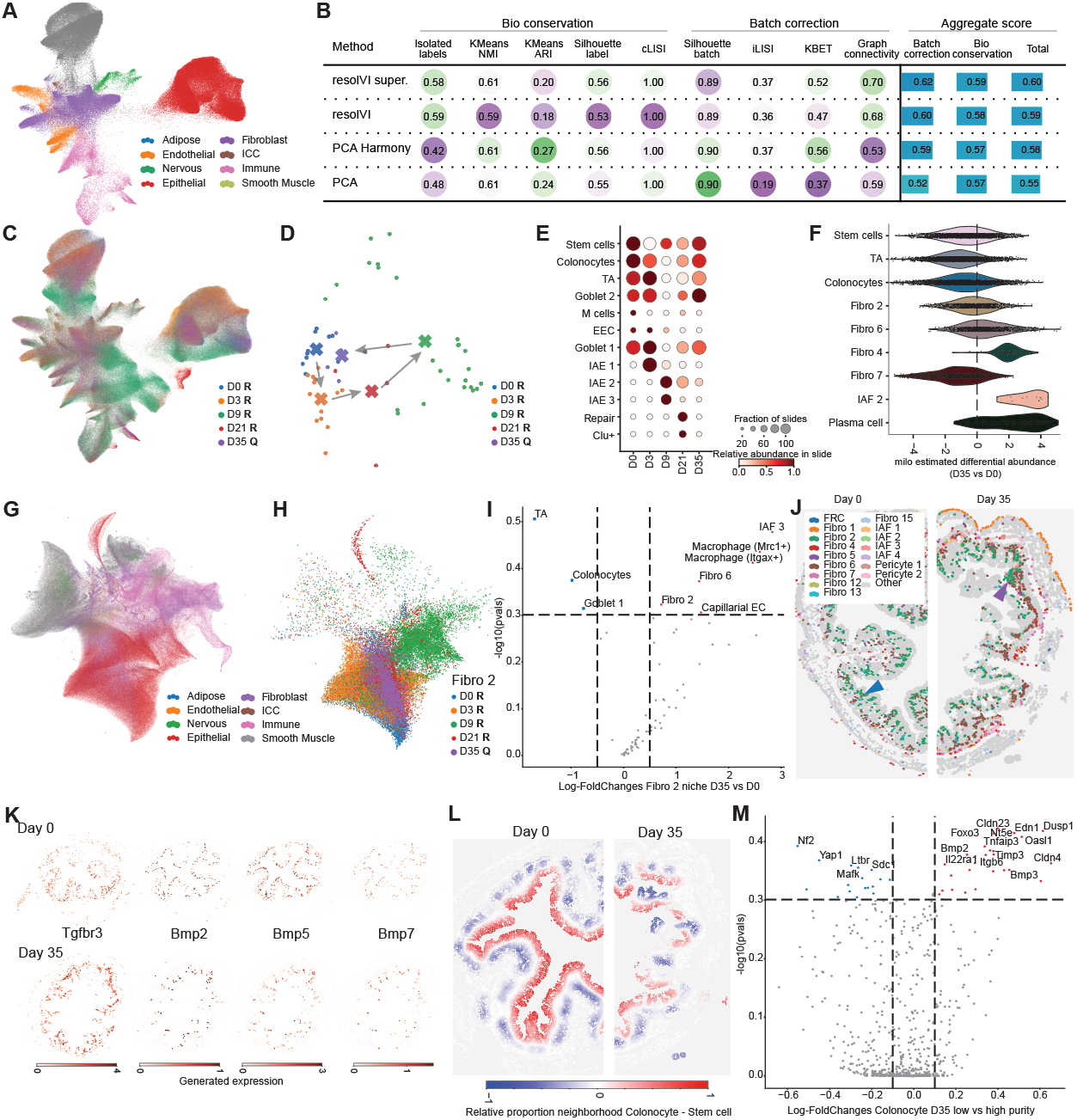
ResolVI reveals alterations in colon epithelium after DSS colitis. **A** UMAP after integration with semi-supervised resolVI and subset to only cells at day 0 colored by coarse cell-types. **B** Scib-metrics using slice ID as the batch key and the original fine cell-type labels as cell type label. Data was subset to day 0 prior to computing metrics and low quality cells were removed. We subset to day zero as we expect all biopsies at day zero to be similar, while at later time points disease induces variation within the same cell-type. **C** UMAP colored by timepoints after DSS. Day 35 was used as a query dataset (Q) and was co-embedded with the cells from other time points (R). **D** We compute the frequency of each cell-type within each slice. Displayed is a PCA plot of these slice-level values of cell-type frequency. We highlight the average embedding at each time point with crosses and show the distinct slices with dots. We find samples at day 9 contain vastly different cell-types, whereas samples at day 35 are similar to day zero. **E** Relative cell-type frequency of epithelial cell types at the various time points. We compute the average epithelial cell-type abundance in each slide and summarize the results by time points. Dot size represents the number of slices with a relative abundance greater than 0.01 and color is the per cell-type standardized abundance. Especially samples at day nine and day 21 contain distinct inflammation associated cell-types not present in the healthy colon. **F** Milo was used to compute neighborhood differential abundance between day zero and day 35. We highlight here a manual selection of cell-types that corrobarates the findings in the original manuscript (plasma cells, IAF2, Fibro 4). In addition, we find a tendency towards reduced epithelial cells, while for Fibro 2 (crypt tip fibroblast) we find no change in cell-type abundance. **G** The cell-type frequency across the 30 nearest neighbors of each cell are computed. This yields for every cell a vector of neighborhood abundances across all cell-types. These vectors are embedded using UMAP. We highlight the cells based on their coarse label. It becomes evident that there exist a neighborhood of epithelial cells, smooth muscle cells, fibroblasts and immune cells. **H** Neighborhood composition UMAP subset to Fibro 2 cells colored by time points. It becomes evident that these cells colocalize with distinct epithelial niches at different time points. **I** We compute relative changes in cell-type composition in the neighborhood of Fibro 2 cells using resolVI to compare fibroblasts at day 35 and day zero. Displayed is the log-2 fold change between cells from both time points and the p-value. **J** Spatial distribution of fibroblasts at day zero (left) and day 35 (right). We find fibroblasts at day zero interspersed with colonocytes (blue arrow), while these cells colocalize with other fibroblasts in aggregates of fibroblasts in the submucosa at day 35 (purple arrow). **K** Comparison of resolVI generated gene expression of fibroblasts at day zero (top row) and day 35 (bottom row). We show *Tgfbr3* as an example of a gene upregulated in proximity to stem cells and higher expressed at day 35 and *Bmp2, Bmp5* and *Bmp7* all higher expressed close to colonocytes and at day zero. **L** Same biopsies as in **J** colored by difference in relative frequency of colonocytes and stem cells in their neighborhood (30 nearest neighbors). **M** Differential expression computed at day 35 between colonocytes with a relative proportion of colonocytes in their neighborhood above 0.5 compared to colonocytes with fewer colonocytes in their neighborhood. The computation was performed using the resolVI differential expression function.

Representing all time points according to their respective cell type frequencies (Figure 5D), we found the highest deviation from the pre-disease tissue nine days after DSS challenge. The lowest deviation is observed at the recovery stage, 35 days after DSS. However, some differences persist even at this late time point. Within epithelial cells, we find a shift towards inflammation-associated epithelial subset 1 (IAE1) on day three, subsets 2, 3 (IAE2/3) on day nine and repair-associated epithelial cells at day 21 (Figure 5E). At day 35 we find more goblet 2 cells but fewer transitional amplifying (TA) cells. To quantify this, we performed a differential cell-type abundance test, applying Milo [26] to the resolVI embedding of the day zero (pre-disease) vs. day 35 samples. We found a decrease on day 35 of stem cells, colonocytes, and TA cells. In addition, we confirmed the findings of the original study, which reported an increase on day 35 of plasma cells, fibroblast 4 and inflammation-associated fibroblast 2 (IAF 2), (Figure 5E).

To study changes in their spatial context, we next considered the composition of small tissue areas, rather than individual cells. To this end. we represented each cell by the vector of predicted cell-type frequencies among its nearest 30 spatial neighbors (Methods, Supplementary Figure 24). The embedding of these representations in UMAP suggests a marked rewiring in the composition of the neighborhoods between day zero and day 35, especially around type 2 fibroblasts, which were interpreted in the original study as tip-crypt fibroblasts (Figure 5G and H). Specifically, we found on day 35 a decrease in co-localization of type 2 fibroblasts with transient amplifying epithelial cells (TA), colonocytes, and goblet 1 cells using a differential colocalization test within resolVI (Methods, Figure 5I). In contrast, we find an increased co-localization of type 2 fibroblasts with stromal cells, such as IAF 3, Fibro 6 and 2 and different macrophages (Figure 5I). Considering the spatial organization of Fibro 2, we find these cells on day zero interspersed between colonic crypts reaching the top of each crypt, while on day 35 we find dense aggregates of Fibro 2 cells in the submucosa (Figure 5J). To study gene expression changes, we first identified markers of Fibro 2 cells colocalizing with colonocytes by computing differentially expressed genes those cells co-localizing with stem cells and those co-localizing with colonocytes (tip crypt; Supplementary Figure 25). We find an increase in the expression of several *Bmp* genes, which are essential for the normal development of colonocytes along the crypt axis [27].

In agreement with a loss of Fibro 2 cells that co-localize with colonocytes, we also find a decreased expression of Bmp genes in Fibro 2 cells on day 35 (Figure 5K). Corroborating a disruption in epithelial cell maturation, while we find a distinct gradient of stem cell to colonocyte transition on day zero along the crypt axis, on day 35, we find an overall disruption in the organization of the epithelial layer (Methods, Figure 5L). To study these disruptions, we performed a differential expression analysis between colonocytes in areas of high colonocyte density and other areas on day 35. We discover higher expression in regions with many colonocytes of barrier-promoting genes like *Cldn4, Cldn23, Tnfaip3* and *Il22ra1* (Figure 5M, Supplementary Figure 26. This postulates a loss of barrier function even long after acute DSS inflammation and after macroscopic regeneration has ensued. Increased barrier permeability is a known issue in inflammatory bowel disease even after control of acute inflammation [28].

## 3 Discussion

We have developed resolVI, a probabilistic modeling approach to clean noise and bias in spatial transcriptomics data analysis. Analysis of spatial transcriptomics data sets on the cell level requires segmentation of cells. Wrong segmentation of cells assigns molecules to neighboring cells leading to spurious coexpression patterns. We demonstrate here that in various segmentation approaches that depend on image or molecule position, spurious coexpression exists, and that resolVI reduces this coexpression pattern across all tested technologies and segmentation algorithms. While resolVI reduces bias across all segmentation, the performance of resolVI depends on the quality of the cell segmentation, and we found the lowest amount of false co-expression using Baysor segmentation. In practice, for large data sets, we recommend initial segmentation using ProSeg as it is more scalable than Baysor.

We show improved cell-type resolution, integration, and downstream analysis, such as differential expression between different spatial niches and differential neighborhood composition analysis, using resolVI.

Applying resolVI to a study of human liver cancer, we demonstrated heterogeneous spatial niches with a pro-inflammatory and anti-inflammatory phenotype of cancer cells and corresponding changes in the phenotype of myeloid and T cells and demonstrated that resolVI generated expression preserves gene expression differences between tumor and healthy tissue.

To highlight resolVI’s downstream capabilities, we applied resolVI to a longitudinal analysis of DSS colitis. We demonstrated that resolVI can efficiently map query samples to an existing reference dataset. To further improve integration and perform label transfer, we provide a semi-supervised mode in resolVI that provides uncertainties for cell-type prediction. Using these uncertainties, we provide a spatial niche embedding of spatial neighborhoods and a framework for differential abundance testing of cell-types for the spatial neighborhood. Using this framework, we found a change in the location of the crypt-tip fibroblasts 35 days after DSS colitis. At this point, we would expect a restoration of a healthy colon. However, these fibroblasts localize in healthy mice in the tip of the crypt and provide Bmp signaling to enterocytes, while after DSS colitis these cells co-localize with other fibroblasts in aggregates in the submucosa. In line with reduced Bmp signaling, we demonstrate a reduced frequency of barrier-promoting epithelial cells. Spatial transcriptomics of human IBD samples will reveal the translational relevance of these findings.

ResolVI is part of the scvi-tools framework and is currently provided at https://github.com/scverse/scvi-tools/pull/3144.

## Acknowledgements

We thank Florian Ingelfinger and Nathan Levy for the early adoption of resolVI and critical feedback. We acknowledge the members of the Yosef laboratory for general feedback. C.E. was supported by the German Research Council (DFG Walter Benjamin Stipendium 448802458).

## Author contributions

C.E. and N.Y. conceptualized the study. C.E. designed and implemented resolVI and performed all experiments. C.E. and N.Y. wrote the manuscript.

## 4 Methods

resolVI is a Python package and is implemented in scvi-tools. resolVI is available within scvi-tools and as such integrates into Python and R workflows. Further details on resolVI and the source code are available at https://github.com/scverse/scvi-tools/pull/3144 and a tutorial is available at https://github.com/scverse/scvi-tutorials/pull/392.

### Model

ResolVI assumes that the observed expression *x*_*ng*_ of the cell *n* and the gene *g* is sampled from an underlying state defined by the true expression of the cell (*z*_*n*_), the gene expression of neighboring cells *N* (*n*) that were incorrectly segmented, and the background gene expression *bg*_*g*_. The generative model of resolVI writes as:

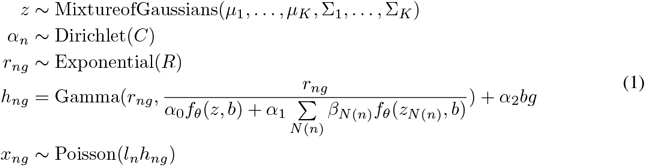

In particular, *z* and *z*_*N*(*n*)_ are the latent embeddings of the cell itself as well as its spatial neighbors both of dimension *L*. ResolVI uses a mixture of Gaussians prior on *z*:

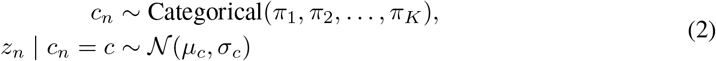

In practice, we assume the covariance matrices to be diagonal and learn *µ*_1_,…, *µ*_*K*_, *σ*_1_,…, *σ*_*K*_, and *π*_1_,…, *π*_*K*_ during training using maximum likelihood estimation.

*α*_*n*0…2_ are the mixture proportions learned via a diffusion encoder which takes as input the gene expression of the cell itself *x*_*ng*_ and encodes this, *α*_*n*_ = *g*(*x*_*ng*_, *b*) followed by a softmax output function. We perform maximum a posteriori (MAP) sampling for *α*_*n*0 2_. The prior C is a hyperparameter that we learn using a Gamma distribution prior. Throughout the manuscript, we set the prior concentration for the true proportions to 3 and the prior for diffusion to 2 and the rate to the concentration divided by 20. For the prior of the background proportion *B*, we expect it to be similar to the counts of negative spike-ins. As these are missing for some technologies, we empirically found that the measured counts of negative spike-ins are similar to the counts of the lowest expressed genes for Nanostring CosMx. We take advantage of this observation to estimate the background and set it to the mean expression of the lowest expressed genes (by default the lowest 10%).

While we assume a Poisson distribution of the background expression, for true expression we use a negative binomial distribution. *r*_*ng*_ is the rate of the negative binomial distribution. To combine both distributions, we use the Gamma-Poisson formulation of the negative binomial distribution. We use an exponential prior for 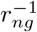 that encourages small values (leading to a Poisson distribution) with *R* = 1 throughout the manuscript.

*f*_*θ*_ is the decoder function shared between the cell *n* and all neighboring cells.

*β*_*nN*(*n*)_ are the contributions of each neighbor (by default 20 neighbors) to the expression caused by diffusion. We use MAP inference for these parameters, and it is not amortized (matrix of *n*_*cells*_*Xn*_*neighbors*_). We assume that cells that are close have a higher likelihood to contribute to wrong segmentation. We use a Radial-Basis-Function kernel to incorporate this into a Dirichlet prior: *γ*_*n*_ ∼ Dirichlet(*exp*(*SD*_*N*(*n*)_*/D*)), with *D*_*N*(*n*)_ being the Euclidean distance between a cell *n* and the respective neighbor, a normalization factor *D*\* by default set to the median distance across all cells to the fifth closest neighbor, and S being a learnable scale parameter (*S* Exponential(1)). As background, we use a prior *bg*_*sg*_ ∼ Dirichlet(1) and learn it separately for each batch *s*.

Taking together, *α*_0_…_2_ adds up to one and the learned background expression, diffusion component, and true expression each add up to one; therefore, the sum of *h*_*ng*_ is one. We restrict the generative process to a fixed count per cell (analogous to library size in sequencing) *l*_*n*_.

### Data augmentation

In spatial transcriptomics, we find a large number of cells with low counts. Encoders tend to not encode the information of these cells correctly. However, in spatial transcriptomics, filtering out low-count cells reduces key insights, such as cell-cell interactions or spatial niche embeddings. Therefore, we chose a low cutoff of 20 molecules per cell for filtering before running resolVI. To increase information for cells with low counts, we downsample the expression *x*_*ng*_ during training using multinomial subsampling before feeding these subsampled counts into the expression encoder. We downsample counts for each cell in a mini-batch to LogNormal(*X*_*median*_, *X*_*std*_) counts with *X*_*median*_ being the median and *X*_*std*_ being the standard deviation of the log-library size in all cells. This encourages the encoder to learn from these cells with few counts. For neighboring cells *N* (*n*) we only apply the learned encoder and don’t perform subsampling but only for the cell *n*.

### Implementation

ResolVI is built within scvi-tools [7] and uses Pyro [29] as a back-end, which allows GPU-accelerated computing. Although all hyperparameters discussed here are exposed, we used the same default settings throughout all case studies and technologies here. The median Euclidean distance to the nearest 5 neighboring cells, the median molecule counts across all cells, and the prior proportion for the unspecific background are inferred from the data set during setting up the data set for the model. A nearest-neighbor graph is built separately for each sample using the scanpy nearest-neighbor computation [30] and scikit-learn _*kneighbors*_*from*_*graph* [31] to assign neighbors. We used 10 latent dimensions for the encoder, an encoder with 2 hidden layers, a dropout in the encoder of 0.05, 128 nodes per hidden layer in the encoder and 32 nodes per hidden layer in the decoder, and are not encoding covariates (neither latent encoder nor mixture proportion encoder), while the decoder uses batch ID as a covariate.

We use a custom guide function to enable amortization. For amortization of *α*, we use the default scVI encoder with a three-dimensional output without dropout and use a softmax activation function for *alpha*.

During training, we do not compute the gradient for *x*_*N*(*n*)*g*_ in *f*_Θ_. Therefore, for neighboring cells, we only apply inference, while for the cell *n* the decoder is trained. This improves stability and training speed. We use a softmax activation function for the decoder output.

Each resolVI model contains five distinct models: (1) model_unconditioned has no observed gene expression, (2) the generic model has observed gene expression, (3) model_corrected sets the background and diffusion proportion to zero and true proportion to one, and therefore generates unbiased gene expression, (4) model_residual sets the true proportion to zero to yield an estimate of diffusion and background, (5) model_simplified blocks most variables except the estimated counts and the proportions to speed up differential expression computation. Generated counts in the manuscript denote the output of *x*_*ng*_ of model_corrected, while we also implemented corrected counts computed by 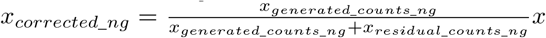 and these counts don’t impute expression.

ResolVI has a combination of amortized and non-amortized parameters, since the contributions to the diffusion expression from each of the neighboring cells *x*_*N*(*n*)*g*_ cannot be amortized. We found empirically that the model during training ignored this parameter as the updates were too infrequent. We fixed this by increasing the learning rate of this parameter to 0.05 and also applied this increased learning rate for the means of the MoG distribution while all other parameters are trained with a learning rate of 1e-3. Unlike scVI, we perform kl_warmup over just 20 epochs of training as faster warmup improves correction of gene expression.

### Supervised scenario

ResolVI takes as optional input prior knowledge about cell type assignment of each cell. We use these inputs in two ways: First, we enforce the mixture of Gaussians to align with cell-type annotations in the supervised scenario. In this case, we set *K* to be the number of cell types and adopt the prior as: *c*_*n*_ ∼ Categorical 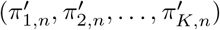, where 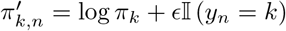 with *ϵ* = 10 by default and *y*_*n*_ the cell type annotation of cell *n* [10].

Second, we train a classifier on latent embedding *z*_*n*_ while training the model that predicts the cell-type label. This encourages distinct cell types to be separated in latent space and additionally allows us to predict cell-type labels. We use a shallow linear classifier in latent space without hidden layers and use 10 as weighting of the loss of the classifier. This yields improved performance to that of the classifier in scANVI.

### Transfer learning

For transfer learning, we adopt the approach from scArches for Pyro models. As some parameters are non-amortized, we define the cell indices as a categorical field and extend this field when performing transfer learning. It is therefore essential that cell indices are unique between query and reference data sets.

For transfer learning, we find the parameters that require updates in scArches (batch-dependent as well as non-amortized parameters) and block the parameters for all other parameters. In addition, we set the learning rate for these parameters to zero. After these adjustments, the model is trained using the same training objective as for the full model.

### Differential expression

We adapt the approach from [10] to Pyro. Specifically, we change the way importance sampling is performed to be in line with Pyro. We use a simplified model in which the loss is restricted to the KL in latent space, the reconstruction and the mixture proportion log-probability. We then perform importance sampling and compute the empirical marginal over the predicted true expression. To encourage sampling cells with low artifacts, we perform weighted sampling weighted by the probability of the observed gene expression given the generated true gene expression. This increases the weight of cells that are well segmented. We divide the log-probability by the counts per cell to avoid preferential sampling of cells with only few counts (reconstruction loss scales with the counts per cell). We plug this estimator into the scvi-tools differential expression function. In short, we compare pairwise samples of gene expression between both groups. The reported probability of DE is the fraction of these samples above a predefined delta. To avoid reporting genes that are not expressed (low likelihood of being expressed), we add a pseudocount to the estimated rates that serves to moderate the pairwise log-2 fold changes. For the summary statistic log-2 fold change, we compute the values without adding the pseudocount to not bias this estimate.

### Differential niche abundance

The calculation is similar to the differential expression function. Instead of performing importance sampling, we take random samples from the latent code of all neighboring cells and predict the cell-type probability using the cell-type classifier. To yield a vector of length number of cell types for each cell, we weight the vector based on the spatial distance to the center cell and sum over all neighboring cells. We perform pairwise comparison for each cell type instead of each gene in the differential expression function.

### Niche embedding

We compute the weighted (by spatial connectivity) average of the cell-type predictions in the spatial neighborhood of each cell. These cell-type frequency vectors are embedded using UMAP.

### Segmentation

For the brain 10X Xenium data, we use xeniumresegment and perform nuclei segmentation by setting the expansion distance to zero. For Baysor, we set the prior segmentation to the original cell segmentation with a confidence of 0.2 and run the segmentation after subsetting the tissue to 70 regions. We set the minimum molecules per cell to 30 and performed segmentation in three dimensions. For ProSeg, we used the nucleus segmentation as the prior segmentation and used the defaults ProSeg settings by setting the Xenium flag.

For Vizgen liver cancer data, we use the vizgen post-processing tool https://github.com/Vizgen/vizgen-postprocessing. We use their example algorithm using Cellpose with one and three z layers and with nuclei segmentation across three z layers. For Baysor, we used the three-z-layer cell segmentation as the prior segmentation. We split the data into 300 spatial regions and performed segmentation for each region separately. We perform three-dimensional segmentation and set the minimum molecules per cell to 20. For ProSeg, we used the three-z-layer nucleus segmentations as prior segmentation and used the defaults by setting the MERSCOPE flag. For Nanostring liver data, we use the original segmentation as prior for Baysor and split the data into several hundred spatial regions. We performed segmentation separately.

### Metrics

#### Integration metrics

We use scib-metrics using default values to benchmark integration. We disabled PCR comparison as we expect that a majority of the variation is due to wrong segmentation. Computing this metric is therefore misleading after correction for artifacts.

#### Double positive metrics

We use single-cell reference sequencing datasets. We pose that genes not co-expressed in single-cell sequencing but co-expressed in spatial transcriptomics are likely due to technical issues and discuss other cases separately. This approach is derived from [32]. We used a Poisson mixture model with two components (foreground and background). We don’t use zero as the threshold as we argue that if fore- and background can be clearly distinguished, it is easy to define a threshold that separates true and false positives. Using this approach applied to the reference dataset, we selected all genes expressed in more than 5% of the cells in one cell type and less than 1% in all other cell types for the brain dataset. For the single nuclei sequencing dataset from the liver, we found more co-expression of marker genes likely due to ambient expression in this single nuclei dataset and selected genes that were expressed in more than 20% of cells in one cell type and less than 10% in all other cell types. For each case, we provide the estimated double positives in the reference dataset as a Supplementary Figure. To evaluate the spatial dataset, we first used the same approach with a Poisson mixture model and computed for each pair of genes the ratio of double positives divided by the number of cells that are at least single positive. In addition, we computed a cosine similarity score between the summed expression of all marker genes of two cell types. For both metrics, we performed count normalization before computing these metrics.

For the 10X Xenium brain data, we yielded the following marker genes: “Vascular”: *Adgrl4, Cldn5, Emcn, Nostrin, Pln, Slfn5, Sox17*; “Neurons”: *Bcl11b, Cabp7, Cbln1, Cbln4, Chrm2, Cntnap4, Cpne4, Fibcd1, Gsg1l, Hs3st2, Lamp5, Ndst4, Necab1, Nell1, Neurod6, Nwd2, Plcxd3, Rxfp1, Satb2, Slc17a6, Sncg, Syt2, Syt6*, “Immune”: *Cd53, Ikzf1, Lyz2, Siglech, Spi1, Trem2*, “Oligodendrocyte”: *Sema3d*, “Ependymal”: *Spag16, Trp73*. We manually added *Gjc3* for oligodendrocyte in Figure 1.

For the Vizgen MERSCOPE data, we yielded the following marker genes: “Fibroblast”: *ACTA2, ANGPT1, COL1A1, COL5A1, COL6A3, HGF, PDGFRA, VEGFC*, “Bcell”: *BLK, CD79B, DERL3, FCRL5, MZB1, POU2AF1, SRPRB, STAP1, TNFRSF13B*, “Myeloid”: *CD163, CD86, CMKLR1, CSF1R, CSF3R, FCGR2A, FCGR3A, HLA-DRB1, IL1R2, ITGAM, LYZ, MAFB, MARCO, MSR1, SIGLEC1, TGFBI, TLR2*, “Lymphocyte”: *CD2, ITK, KLRF1, KLRG1, PRF1*, “Endothelial”: *CDH5, CLDN5, FLT1, FLT4, KDR, KLF2, MMRN1, NOS3, TEAD4*.

For the Nanostring CosMx data, we yielded the following marker genes: “Myeloid”: *ADGRE2, C1QA, C1QB, C1QC, CD163, CD68, CD86, CLEC7A, CMKLR1, CSF1R, CSF3R, FPR1, GPNMB, HCK, IL1R2, LYZ, MARCO, MS4A4A, MSR1, TLR2*, “Endothelial”: *ADGRL4, CD9, CLEC1A, FLT1, FZD4, IL33, KDR, NPR1, RAMP3, TIE1*, “Fibroblast”: *ANGPT1, BGN, BMP5, CACNA1C, CARMN, CDH19, COL12A1, COL14A1, COL1A1, COL1A2, COL3A1, COL6A3, EPHA3, HGF, IGFBP5, MYL9, PDGFRA, RAMP1, VEGFC*, “Cholangiocyte”: *CASR, CCL28, CD24, IL20RA, ITGA2, ITGB8, KRT7, SPP1*, “Lympho”: *CCL5, CD2, IL7R, ITK, KLRF1, PRF1*, “Bcell”: *CD27, IGHA1, IGHG1, IGHM, IGKC, JCHAIN, MZB1, SELL, TNFRSF13B, WNT5B*.

### Downstream analysis

Throughout the manuscript, we use generated expression from resolVI. We perform count normalization. Recently, area normalization was suggested for spatial transcriptomics [33]. However, this benchmarking was performed without comparing different segmentation algorithms. We perform square root transformation of normalized counts as we found after log-1p transformation a worse identification of expected cell types. For all UMAP plots, we first compute 20 nearest neighbors using RAPIDS and then perform UMAP using the implementation in the UMAP package [34].

For all plots, we first perform cell typing in one of the segmentations. For 10X Xenium and Vizgen MERSCOPE, we performed Leiden clustering and manual annotation after ProSeg segmentation on ProSeg estimated counts. For Nanostring CosMx and colitis data, we used the original cell types provided by the authors.

We then trained a supervised resolVI model on this segmentation. To transfer labels to other segmentations, we used these other segmentations as query data sets of this reference model and transferred labels to the other segmentations. Especially for Nanostring CosMx data, we found a tendency to label many cells as hepatocytes and tumor cells that have a mixed phenotype between immune cells and epithelial cells. We recomputed cell types using Leiden clustering after performing resolVI correction and decided to not compute integration metrics on this data set. Leiden clustering was performed with manually selected resolution to capture manually identified differences in gene expression.

### Liver Nanostring CosMx analysis

In panel A, we highlight the count-normalized gene expression of *SAA1* minus the expression of *GLUL* this highlights the peri-central to -portal gradient. We calculate the hotspot modules in the generated expression using the spatial position as the embedding space. In panel H, we compute the differential expression using resolVI between cells positive for the anti- and pro-inflammatory hotspot module. For macrophage and T cell analysis, the Leiden clustering resolution was manually selected. Cell types were manually labeled.

### Colitis data

We first trained an unsupervised resolVI model in cells from day zero to 21. We perform a *k* NN classification of cells in the latent space of resolVI. To this end, we compute a *k* NN graph with 100 nearest neighbors using scanpy and use the neighborhood connectivities as weights for the classifier. The certainty of prediction is the sum of connectivities with that label divided by the sum over all connectivities. We set the label to unknown for all cells with an uncertainty above 5% for their coarse label (Tier1) and above 40% or more for their fine label (Tier3). After removing the label of these cells, we train a semi-supervised resolVI model on the fine labels and predict the label of all cells. We use these labels throughout the rest of the figure. To integrate data from day 35, we performed query mapping using scArches for these cells.

In panel D, we compute the average cell-type frequency across the full slides and embed this frequency vector using principal component analysis. In panel F, we performed milo analysis between cells from time point day 0 and day 35. We used 100 nearest neighbors, proportion 0.1 and used the time point as the design without any covariates. To compute the violin plots, we use the annotate_nhoods function and set neighborhoods with a cell-type fraction below 0.4 to Mixed.

For panel G, we first compute the 30 nearest neighbors using the squidpy spatial_neighbors function using the slice as the library key with Delaunay set to false. We compute the weighted average over the predicted cell-type labels in the spatial neighborhood using the neighbor connectivity as weight. After performing this weighted average, we compute 50 nearest neighbors on these cell-type frequency vectors and compute a UMAP embedding. In panel I, we use the differential abundance function in resolVI comparing predicted fibroblast cells at time point day 0 within the MU3 (stem cell) niche and MU1 (colonocyte) niche. We use the differential expression function in resolVI using a pseudocount of 0.01, delta of 0.05, and filter outlier cells.

To compute the gradient in panel L, we compute the fraction of colonocytes and stem cells in the spatial neighborhood frequency vectors of each cell and subtract the fraction of colonocytes from the fraction of stem cells. To compute differentially expressed genes in colonocytes in panel M, we use the same settings as for fibroblast and compare cells with a difference between colonocytes and stem cells above 0.5 at day 35.

## Code availability

The code to reproduce the experiments of this manuscript is available at https://github.com/YosefLab/resolvi-reproducibility.

## Data availability

The Xenium brain data was downloaded from 10X: https://www.10xgenomics.com/datasets/fresh-frozen-mouse-brain-for-xenium-explorer-demo-1-standard.

The single cell brain reference data set was downloaded from http://mousebrain.org/adolescent/downloads.html.

The Vizgen liver cancer data was downloaded from https://console.cloud.google.com/storage/browser/vz-ffpe-showcase.

The single nuclei liver reference data set was downloaded from https://cellxgene.cziscience.com/collections/0c8a364b-97b5-4cc8-a593-23c38c6f0ac5. Data was subset to all cells from healthy donors.

Nanostring CosMx liver cancer data was downloaded from https://nanostring.com/products/cosmx-spatial-molecular-imager/ffpe-dataset/human-liver-rna-ffpe-dataset/. Colitis data was downloaded from https://datadryad.org/stash/dataset/doi:10.5061/dryad.rjdfn2zh3. We used the h5ad files from day 0-21 and day 35 and the csv files with raw expression values. All processed data will be uploaded to Dryad.

## References

[1] Dario Bressan, Giorgia Battistoni, and Gregory J Hannon. “The dawn of spatial omics”. en. In: Science (New York, N.Y.) (6657 2023). : 10.1126/science.abq4964 (visited on 11/19/2024).

[2] Carsen Stringer, Tim Wang, Michalis Michaelos, and Marius Pachitariu. “Cellpose: a generalist algorithm for cellular segmentation”. en. In: Nature methods (1 2021). : https://www.nature.com/articles/s41592-020-01018-x (visited on 10/04/2024).

[3] Noah F Greenwald, Geneva Miller, Erick Moen, Alex Kong, Adam Kagel, Thomas Dougherty, Christine Camacho Fullaway, Brianna J McIntosh, Ke Xuan Leow, et al. “Whole-cell segmentation of tissue images with human-level performance using large-scale data annotation and deep learning”. en. In: Nature biotechnology (4 2022). : https://www.nature.com/articles/s41587-021-01094-0 (visited on 11/19/2024).

[4] Jun Ma, Ronald Xie, Shamini Ayyadhury, Cheng Ge, Anubha Gupta, Ritu Gupta, Song Gu, Yao Zhang, Gihun Lee, et al. “The multimodality cell segmentation challenge: toward universal solutions”. en. In: Nature methods (6 2024). : https://www.nature.com/articles/s41592-024-02233-6 (visited on 01/03/2025).

[5] Viktor Petukhov, Rosalind J Xu, Ruslan A Soldatov, Paolo Cadinu, Konstantin Khodosevich, Jeffrey R Mott, and Peter V Kharchenko. “Cell segmentation in imaging-based spatial transcriptomics”. en. In: Nature biotechnology (3 2022). : https://www.nature.com/articles/s41587-021-01044-w (visited on 10/04/2024).

[6] Daniel C Jones, Anna E Elz, Azadeh Hadadianpour, Heeju Ryu, David R Glass, and Evan W Newell. “Cell simulation as cell segmentation”. en. In: bioRxiv.org: the preprint server for biology (2024). : https://www.biorxiv.org/content/10.1101/2024.04.25.591218v2.abstract (visited on 10/04/2024).

[7] Adam Gayoso, Romain Lopez, Galen Xing, Pierre Boyeau, Valeh Valiollah Pour Amiri, Justin Hong, Katherine Wu, Michael Jayasuriya, Edouard Mehlman, et al. “A Python library for probabilistic analysis of single-cell omics data”. en. In: Nature biotechnology (2 2022). : 10.1038/s41587-021-01206-w.

[8] Romain Lopez, Jeffrey Regier, Michael B Cole, Michael I Jordan, and Nir Yosef. “Deep generative modeling for single-cell transcriptomics”. en. In: Nature methods (12 2018). : 10.1038/s41592-018-0229-2.

[9] Karin Hrovatin, Amir Ali Moinfar, Luke Zappia, Alejandro Tejada Lapuerta, Ben Lengerich, Manolis Kellis, and Fabian J Theis. “Integrating single-cell RNA-seq datasets with substantial batch effects”. en. In: bioRxiv.org: the preprint server for biology (2024). : https://www.ncbi.nlm.nih.gov/pmc/articles/PMC10635119/ (visited on 10/04/2024).

[10] Pierre Boyeau, Jeffrey Regier, Adam Gayoso, Michael I Jordan, Romain Lopez, and Nir Yosef. “An empirical Bayes method for differential expression analysis of single cells with deep generative models”. en. In: Proceedings of the National Academy of Sciences of the United States of America (21 2023). : 10.1073/pnas.2209124120.

[11] Paolo Cadinu, Kisha N Sivanathan, Aditya Misra, Rosalind J Xu, Davide Mangani, Evan Yang, Joseph M Rone, Katherine Tooley, Yoon-Chul Kye, et al. “Charting the cellular biogeography in colitis reveals fibroblast trajectories and coordinated spatial remodeling”. en. In: Cell (8 2024). : 10.1016/j.cell.2024.03.013.

[12] Malte D Luecken, M Büttner, K Chaichoompu, A Danese, M Interlandi, M F Mueller, D C Strobl, L Zappia, M Dugas, et al. “Benchmarking atlas-level data integration in single-cell genomics”. en. In: Nature methods (1 2021). : https://www.nature.com/articles/s41592-021-01336-8 (visited on 03/13/2023).

[13] Amit Zeisel, Hannah Hochgerner, Peter Lönnerberg, Anna Johnsson, Fatima Memic, Job van der Zwan, Martin Häring, Emelie Braun, Lars E Borm, et al. “Molecular architecture of the mouse nervous system”. en. In: Cell (4 2018). : http://www.cell.com/article/S009286741830789X/abstract (visited on 01/03/2025).

[14] Olga Bondareva, Jesús Rafael Rodríguez-Aguilera, Fabiana Oliveira, Longsheng Liao, Alina Rose, Anubhuti Gupta, Kunal Singh, Florian Geier, Jenny Schuster, et al. “Single-cell profiling of vascular endothelial cells reveals progressive organ-specific vulnerabilities during obesity”. en. In: Nature metabolism (11 2022). : https://www.nature.com/articles/s42255-022-00674-x (visited on 10/07/2024).

[15] MERSCOPE FFPE Solution. en. : https://info.vizgen.com/merscope-ffpe-solution (visited on 01/05/2025).

[16] Ilya Korsunsky, Nghia Millard, Jean Fan, Kamil Slowikowski, Fan Zhang, Kevin Wei, Yuriy Baglaenko, Michael Brenner, Po-Ru Loh, et al. “Fast, sensitive and accurate integration of single-cell data with Harmony”. en. In: Nature methods (12 2019). : 10.1038/s41592-019-0619-0.

[17] Tallulah S Andrews, Diana Nakib, Catia T Perciani, Xue Zhong Ma, Lewis Liu, Erin Winter, Damra Camat, Sai W Chung, Patricia Lumanto, et al. “Single-cell, single-nucleus, and spatial transcriptomics characteriza-tion of the immunological landscape in the healthy and PSC human liver”. en. In: Journal of hepatology (5 2024). : http://www.journal-of-hepatology.eu/article/S0168827824000035/abstract (visited on 11/04/2024).

[18] Y Min, S Ghose, K Boelte, J Li, L Yang, and P C Lin. “C/EBP-Delta regulates VEGF-C autocrine signaling in lymphangiogenesis and metastasis of lung cancer through HIF-1Alpha”. en. In: Oncogene (49 2011). : https://pmc.ncbi.nlm.nih.gov/articles/PMC3175299/ (visited on 10/22/2024).

[19] Quick Start (Cell Bin) - Stereopy. : https://stereopy.readthedocs.io/en/latest/Tutorials/CellBin_Clustering.html (visited on 01/05/2025).

[20] CosMx SMI Human Liver RNA FFPE Dataset. en. 2023. : https://nanostring.com/products/cosmx-spatial-molecular-imager/ffpe-dataset/human-liver-rna-ffpe-dataset/ (visited on 01/05/2025).

[21] Patrick Danaher, Edward Zhao, Zhi Yang, David Ross, Mark Gregory, Zach Reitz, Tae K Kim, Sarah Baxter, Shaun Jackson, et al. “Insitutype: likelihood-based cell typing for single cell spatial transcriptomics”. In: bioRxiv (2022). : https://www.biorxiv.org/content/10.1101/2022.10.19.512902v1.

[22] Liu Xu, Yibing Chen, Lingling Liu, Xinyu Hu, Chengsi He, Yuan Zhou, Xinyi Ding, Minhua Luo, Jiajing Yan, et al. “Tumor-associated macrophage subtypes on cancer immunity along with prognostic analysis and SPP1-mediated interactions between tumor cells and macrophages”. en. In: PLoS genetics (4 2024). : https://pubmed.ncbi.nlm.nih.gov/38648200/ (visited on 01/05/2025).

[23] David DeTomaso and Nir Yosef. “Hotspot identifies informative gene modules across modalities of single-cell genomics”. en. In: Cell systems (5 2021). : https://pubmed.ncbi.nlm.nih.gov/33951459/.

[24] Li Yang, Yanli Pang, and Harold L Moses. “TGF-beta and immune cells: an important regulatory axis in the tumor microenvironment and progression”. en. In: Trends in immunology (6 2010). : https://pmc.ncbi.nlm.nih.gov/articles/PMC2891151/.

[25] Mohammad Lotfollahi, Mohsen Naghipourfar, Malte D Luecken, Matin Khajavi, Maren Büttner, Marco Wagenstetter, Žiga Avsec, Adam Gayoso, Nir Yosef, et al. “Mapping single-cell data to reference atlases by transfer learning”. en. In: Nature biotechnology (1 2021). : https://www.nature.com/articles/s41587-021-01001-7 (visited on 03/13/2023).

[26] Emma Dann, Neil C Henderson, Sarah A Teichmann, Michael D Morgan, and John C Marioni. “Differential abundance testing on single-cell data using k-nearest neighbor graphs”. en. In: Nature biotechnology (2 2022). : 10.1038/s41587-021-01033-z.

[27] Judith Kraiczy, Neil McCarthy, Ermanno Malagola, Guodong Tie, Shariq Madha, Dario Boffelli, Daniel E Wagner, Timothy C Wang, and Ramesh A Shivdasani. “Graded BMP signaling within intestinal crypt architecture directs self-organization of the Wnt-secreting stem cell niche”. en. In: Cell stem cell (4 2023). : http://www.cell.com/article/S1934590923000759/abstract (visited on 01/14/2025).

[28] Timo Rath, Raja Atreya, Julia Bodenschatz, Wolfgang Uter, Carol E Geppert, Francesco Vitali, Sarah Fischer, Maximilian J Waldner, Jean-Frédéric Colombel, et al. “Intestinal barrier healing is superior to endoscopic and histologic remission for predicting major adverse outcomes in inflammatory bowel disease: The prospective ERIca trial”. en. In: Gastroenterology (2 2023). : http://www.gastrojournal.org/article/S0016508522011921/abstract (visited on 01/14/2025).

## References

[29] Bingham Eli, P Chen Jonathan, Jankowiak Martin, Obermeyer Fritz, Pradhan Neeraj, Karaletsos Theofanis, Singh Rohit, Szerlip Paul, Horsfall Paul, et al. “Pyro: Deep universal probabilistic programming”. In: arXiv [cs.LG] (2018). arXiv: 1810.09538 [cs.LG]. : http://arxiv.org/abs/1810.09538 (visited on 10/04/2024).

[30] F Alexander Wolf, Philipp Angerer, and Fabian J Theis. “SCANPY: large-scale single-cell gene expression data analysis”. en. In: Genome biology (1 2018). : 10.1186/s13059-017-1382-0.

[31] Fabian Pedregosa, Gaël Varoquaux, Alexandre Gramfort, Vincent Michel, Bertrand Thirion, Olivier Grisel, Mathieu Blondel, Peter Prettenhofer, Ron Weiss, et al. “Scikit-learn: Machine Learning in Python”. In: Journal of machine learning research: JMLR (85 2011). : http://jmlr.org/papers/v12/pedregosa11a.html (visited on 03/13/2023).

[32] Austin Hartman and Rahul Satija. “Comparative analysis of multiplexed in situ gene expression profiling technologies”. en. In: bioRxiv.org: the preprint server for biology (2024). : https://www.biorxiv.org/content/10.1101/2024.01.11.575135v1.abstract (visited on 10/04/2024).

[33] Lyla Atta, Kalen Clifton, Manjari Anant, Gohta Aihara, and Jean Fan. “Gene count normalization in single-cell imaging-based spatially resolved transcriptomics”. en. In: Genome biology (1 2024). : https://genomebiology.biomedcentral.com/articles/10.1186/s13059-024-03303-w (visited on 11/04/2024).

[34] Leland McInnes, John Healy, and James Melville. “UMAP: Uniform Manifold Approximation and Projection for Dimension Reduction”. In: arXiv [stat.ML] (2018). arXiv: 1802.03426 [stat.ML]. : http://arxiv.org/abs/1802.03426 (visited on 01/14/2025).

