## Supplementary Figures for "ResolVI - addressing noise and bias in spatial transcriptomics"

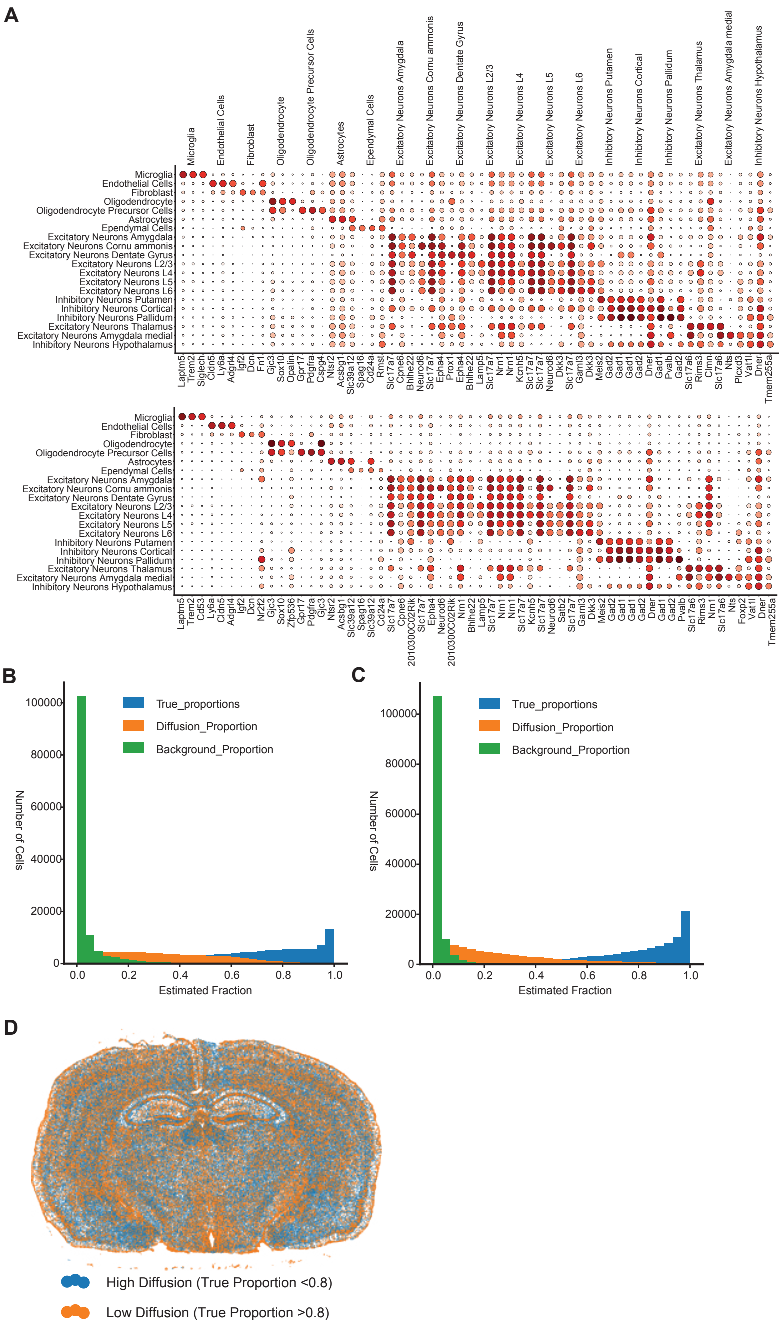


**Supplementary Figure 1: Cell-type definition and estimated true proportion in ProSeg and original segmentation. A** Dotplot of marker genes computed using scanpy default settings displayed for original (top) and ProSeg (bottom) segmentation. Cell-types were defined after ProSeg segmentation and transferred to all other segmentations. Both datasets yield highly similar marker genes. **B** Histogram of diffusion, background and true proportion in the original segmentation. Each entry is one cell. **C** Histogram of diffusion, background and true proportion in the ProSeg segmentation. Each entry is one cell. **D** Spatial display of true proportion in original segmentation. Proportions were threshold at 0.8, as displayed in 2 B. High diffusion lower diffusion is visible in lower cellular density regions in the white matter.

A

| Method | Bio conservation: Isolated labels | Bio conservation: KMeans NMI | Bio conservation: KMeans ARI | Bio conservation: Silhouette label | Bio conservation: cLISI | Batch correction: Silhouette batch | Batch correction: iLISI | Batch correction: KBET | Batch correction: Graph connectivity | Aggregate score: Batch correction | Aggregate score: Bio conservation | Aggregate score: Total |
|---|---|---|---|---|---|---|---|---|---|---|---|---|
| resolVI Supervised | 0.69 | 0.85 | 0.79 | 0.66 | 1.00 | 0.83 | 0.23 | 0.24 | 0.98 | 0.57 | 0.80 | 0.71 |
| PCA Generated Super. | 0.67 | 0.80 | 0.64 | 0.65 | 1.00 | 0.83 | 0.26 | 0.25 | 0.93 | 0.57 | 0.75 | 0.68 |
| PCA Generated | 0.66 | 0.77 | 0.65 | 0.64 | 1.00 | 0.79 | 0.27 | 0.28 | 0.92 | 0.57 | 0.74 | 0.67 |
| resolVI | 0.65 | 0.75 | 0.60 | 0.62 | 1.00 | 0.82 | 0.25 | 0.28 | 0.95 | 0.57 | 0.72 | 0.66 |
| scANVI | 0.65 | 0.79 | 0.67 | 0.62 | 1.00 | 0.87 | 0.11 | 0.18 | 0.98 | 0.53 | 0.75 | 0.66 |
| scVI | 0.60 | 0.71 | 0.59 | 0.57 | 1.00 | 0.90 | 0.14 | 0.21 | 0.97 | 0.56 | 0.70 | 0.64 |
| PCA raw data | 0.60 | 0.75 | 0.61 | 0.58 | 1.00 | 0.92 | 0.16 | 0.19 | 0.83 | 0.52 | 0.71 | 0.63 |

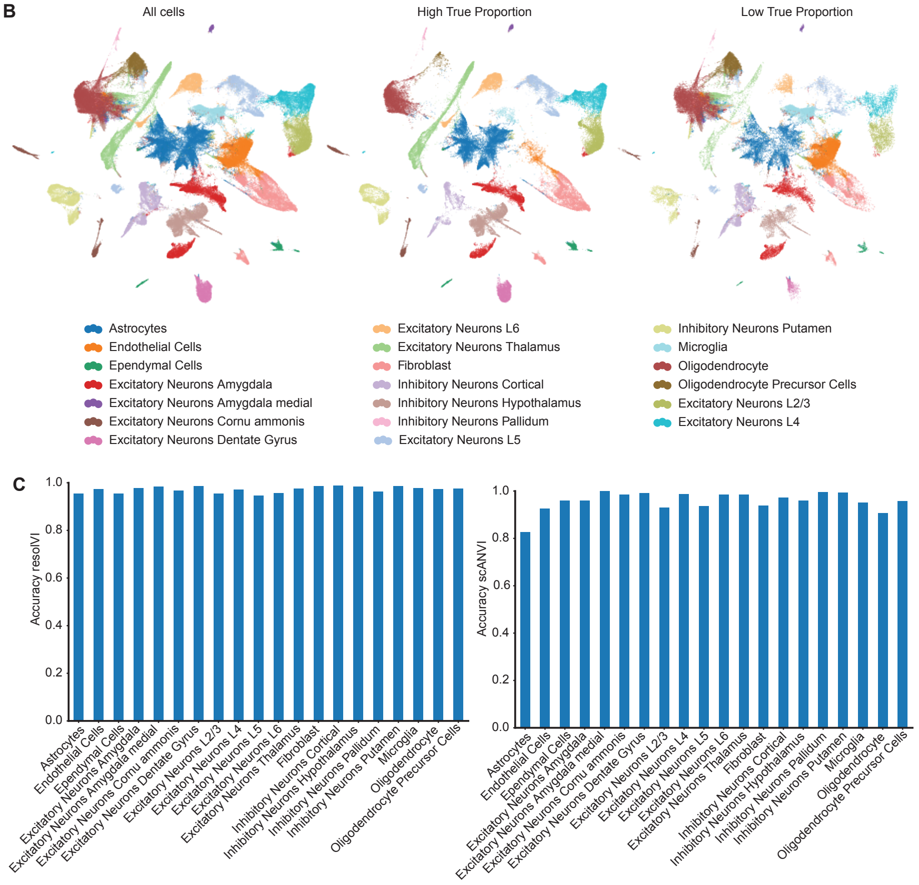


**Supplementary Figure 2: Extended benchmarking of resolVI on Xenium brain data. A** Full scib-metrics scores as described in 2. Generated and Generated Supervised denote PCA computed on the generated true counts estimated by both resolVI models. **B** UMAP computed on resolVI semisupervised model colored by cell-type labels. The right plot shows all cells, the middle plot all cells with an estimated true proportion above 0.8 and the left plot all other cells (batch key in scib-metrics). **C** Both scANVI and resolVI supervised can predict cell-type labels. We benchmark here the per-cell type accuracy of cell-type prediction in both models. For resolVI, accuracy for all cell-types is above 0.95, while accuracy for astrocytes in scANVI is as low as 0.8.

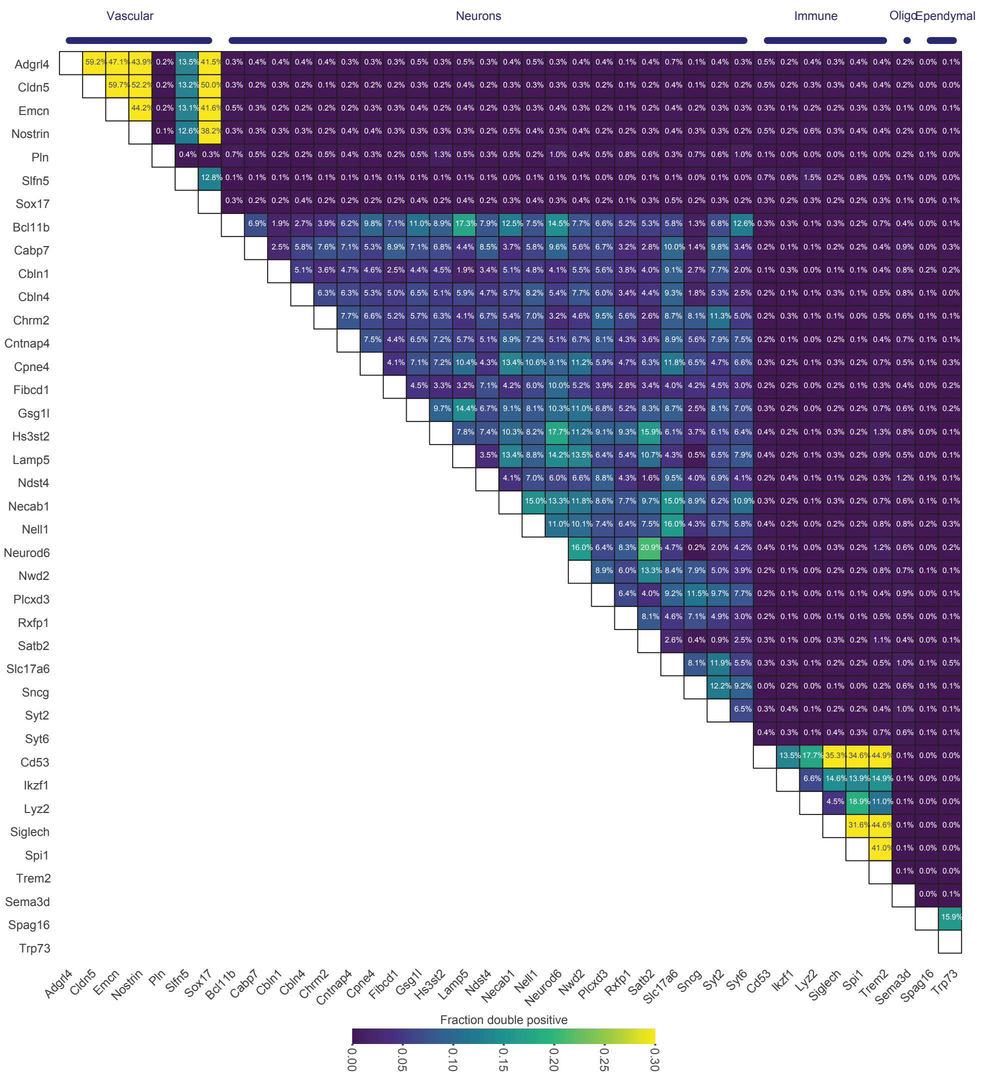


**Supplementary Figure 3: Estimated ratio of double positive expression in single-cell reference dataset.** We subset the single-cell reference dataset to all genes in the Xenium panel and compute a Poisson mixture model on this data. Genes were selected to be expressed in more than 5% of cells of a cell-type and less than 1% in all other cell-types. Displayed is the estimated double positive rate across all cells. The cell-types of the respective genes are highlighted as column annotations. We find co-expression of genes within the cell-types, while genes were not explicitly selected to be co-expressed.

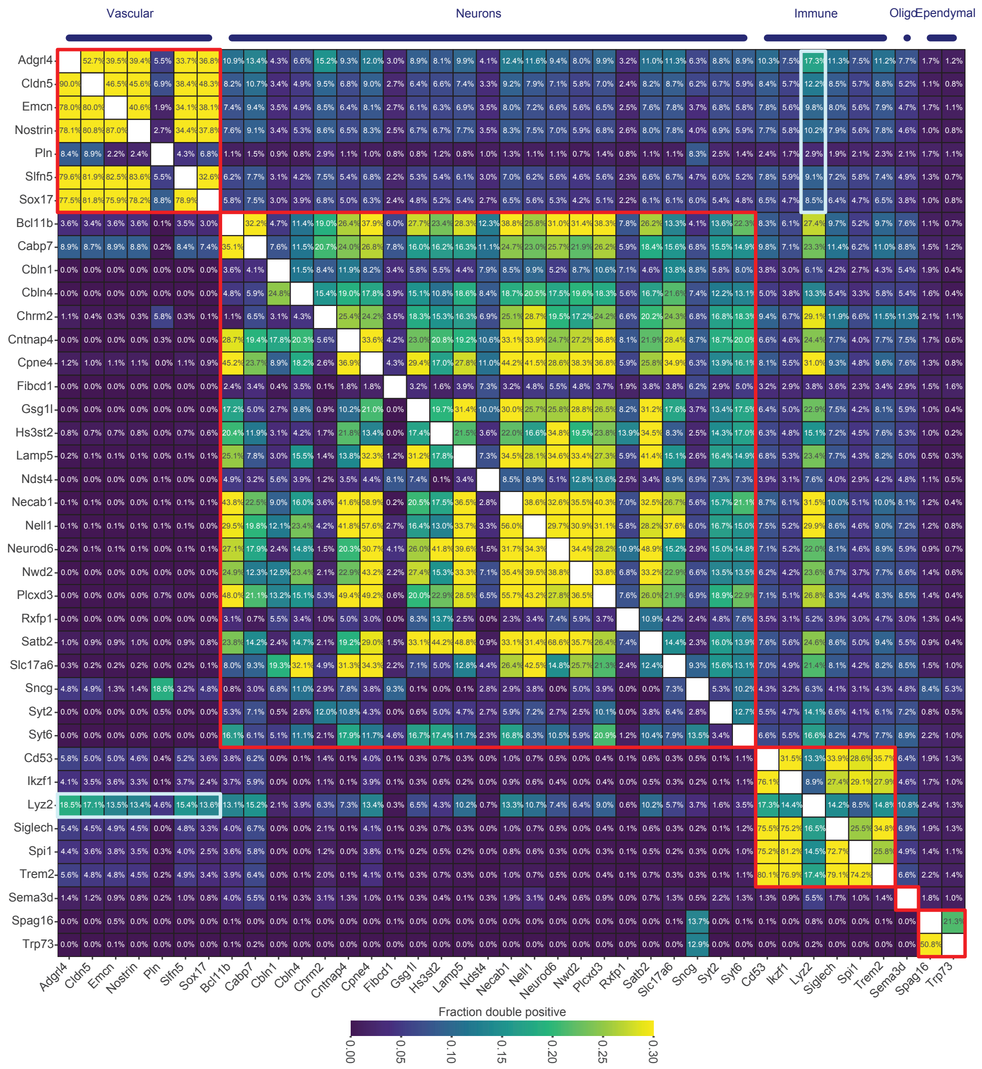

**Supplementary Figure 4: Estimated ratio of double positive expression in Xenium brain data using the original cell segmentation.** We compute a Poisson mixture model on count-normalized data. Displayed is the estimated double positive rate. The cell-types of the respective genes are shown as column annotations. Above the diagonal we highlight expression in raw data, while below the diagonal we highlight resolVI generated data. Red colored boxes show genes that are expressed by the same cell-type (not necessarily co-expressed as they might be cell sub-type specific). We highlight with a turquoise box *Lyz2* co-expression with genes expressed by vascular cells (endothelial cells). This co-expression is preserved by resolVI, while the ratio of double positive of neuron markers and *Lyz2* is reduced by resolVI.

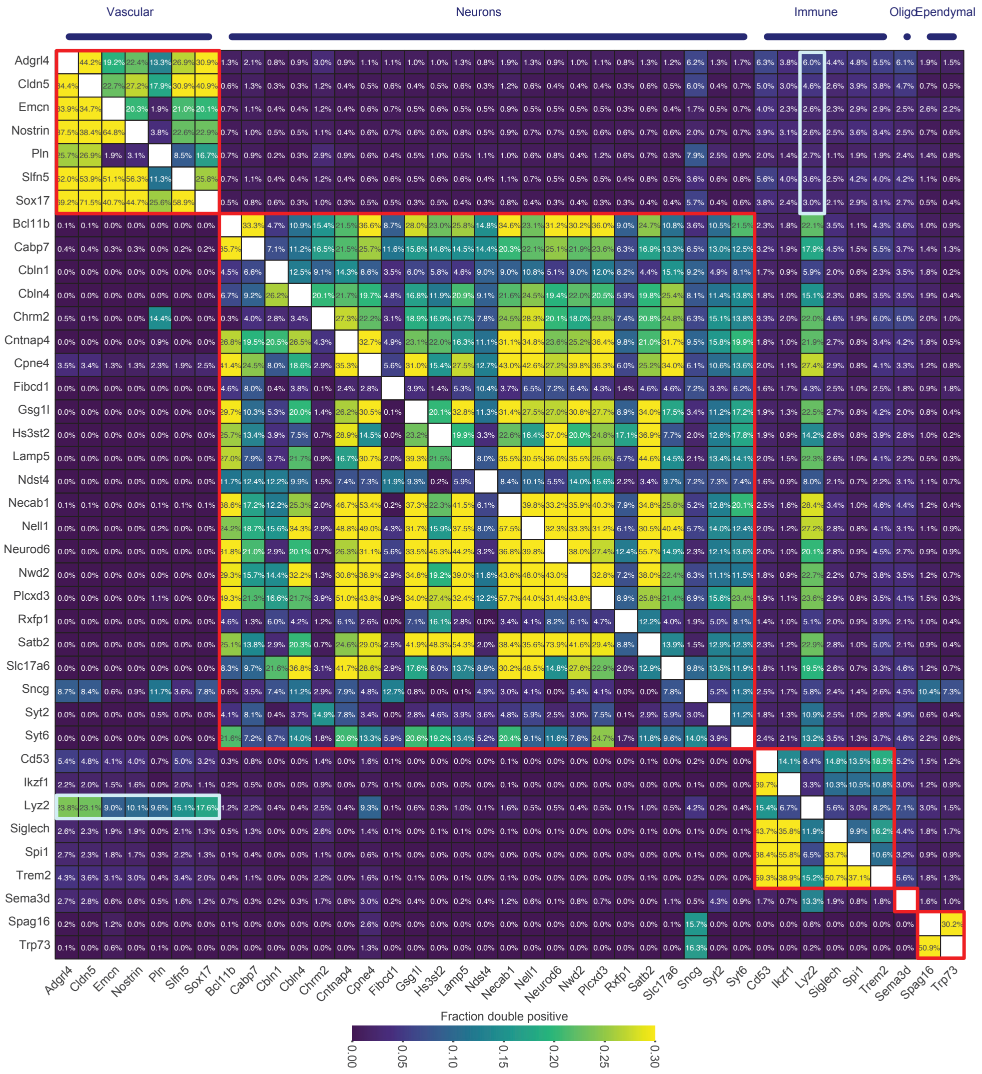


**Supplementary Figure 5: Estimated ratio of double positive expression in Xenium brain data using the original cell segmentation subset to high quality cells.** We computed a Poisson mixture model on count-normalized data. Displayed is the estimated double positive rate. We subset the original data to all cells with an estimated true proportion by resolVI above 0.9 here. These cells are expected to have very little contamination. Displayed is the estimated double positive rate. Above the diagonal we highlight expression in raw data, while below the diagonal fraction of double positives in resolVI generated data is highlighted. In comparison to 4, we see a reduced rate of double positive gene expression before correction.

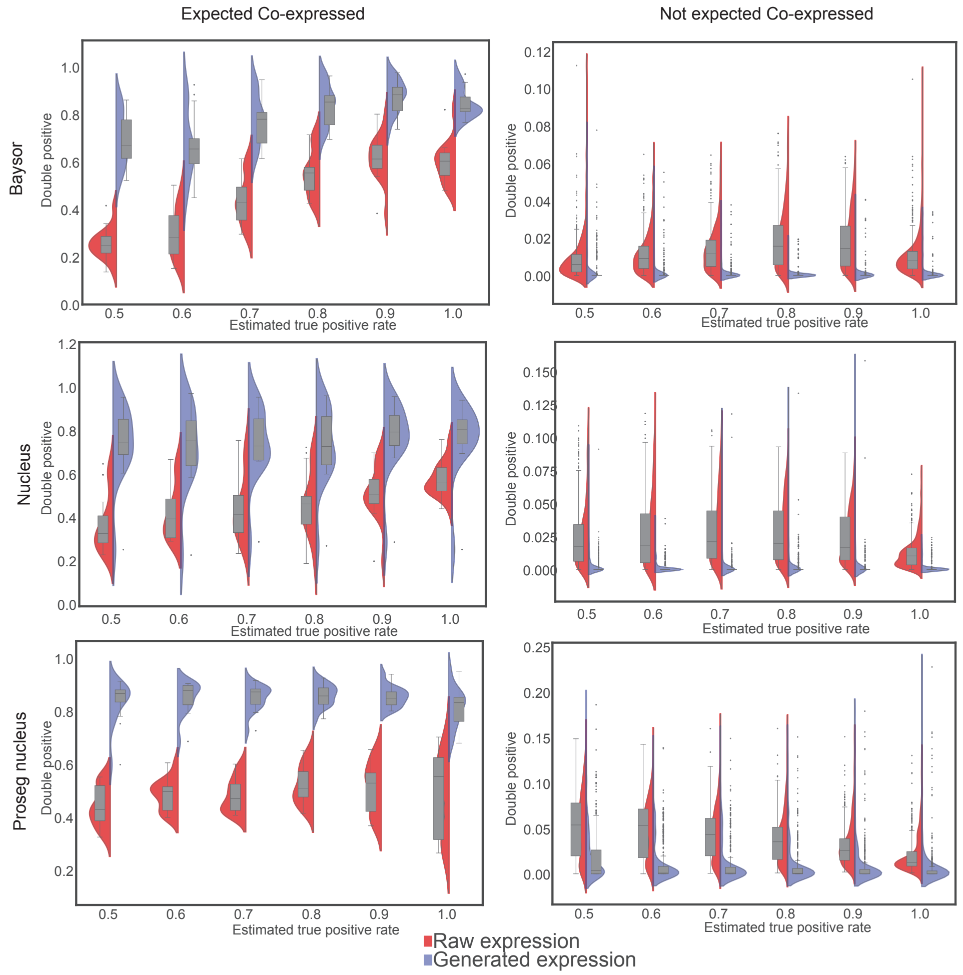


**Supplementary Figure 6: Benchmarking of estimates of true proportions after applying different segmentation algorithms.** The plot is analogue to Figure 2F. In short cells are split into bins with different estimated true proportion. On the left double-positive fraction of genes that are expected to be co-expressed is displayed and on the right pairs of genes that are markers of distinct cell-types are displayed. On the left a higher value is favorable, while on the right lower values are expected. In Figure 2, we used the cell segmentation from 10X Xenium, which performs extension of nuclei segmentation to perform cell segmentation. For all segmentations (Baysor, Xenium nucleus segmentation and ProSeg segmentation) we find improvements over the original count data using generated expression. We find higher double positive rate for co-expressed genes and a lower rate for mutually exclusive expressed genes. Cells with a high true proportion in ProSeg and nucleus segmentation have a lower false double-positive rate before correction.

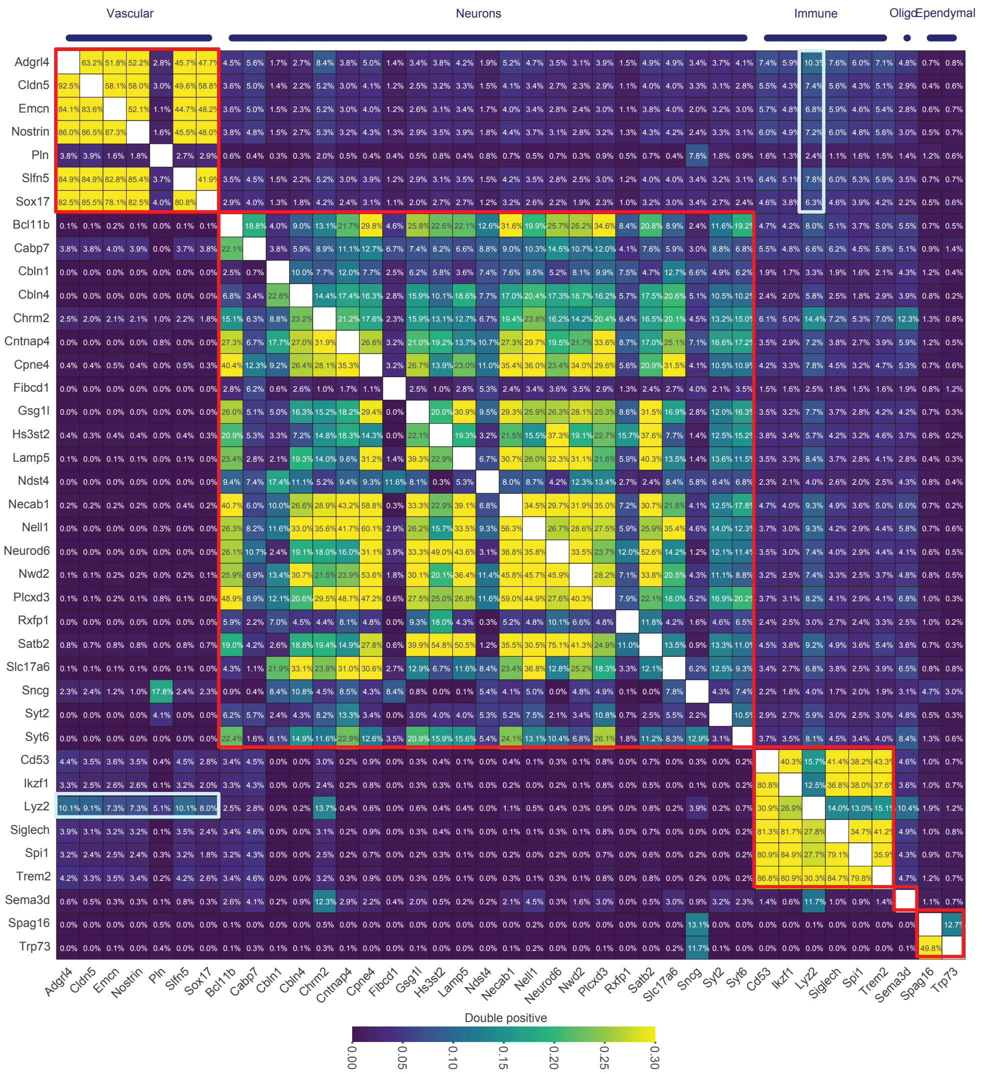


**Supplementary Figure 7: Estimated ratio of double positive expression in Xenium brain data using ProSeg segmentation.** We computed a Poisson mixture model on count-normalized data. Displayed is the estimated double positive rate. Respective cell-types of the marker genes are shown as column annotations. Above the diagonal we highlight expression counts after ProSeg segmentation, while below the diagonal fraction of double positives in resolVI generated data is highlighted. In comparison to 4, we see a reduction of double positive before and after correction. ResolVI predicts again *Lyz2* expression in endothelial cells but not neurons. The outliers of high expression beyond the cell-type for which genes are markers (*Sncg* in ependymal cells and *Sncg*-*Pln* are preserved in both models highlighting agreement across different segmentations.

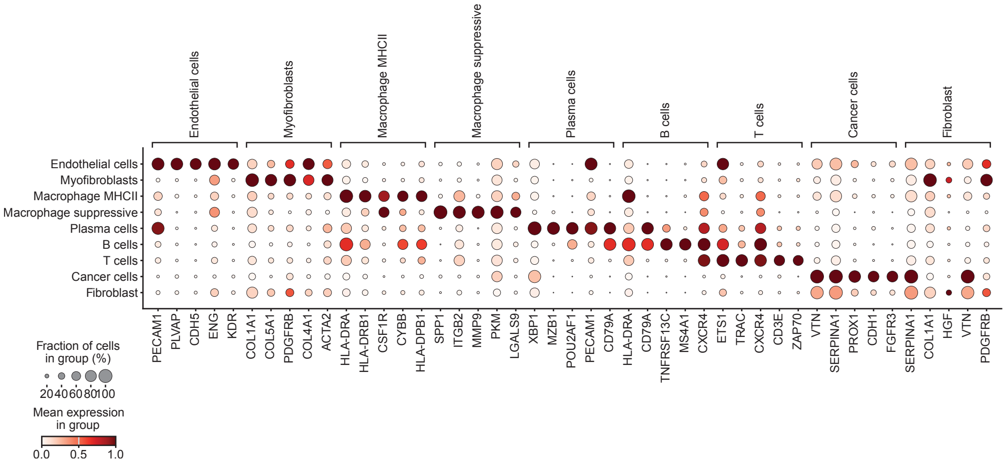


**Supplementary Figure 8: Cell-type definition of Vizgen liver cell-types.** Top-5 marker genes for each cell-type are displayed using scanpy rank gene marker function. Displayed are count-normalized observed counts after ProSeg segmentation.

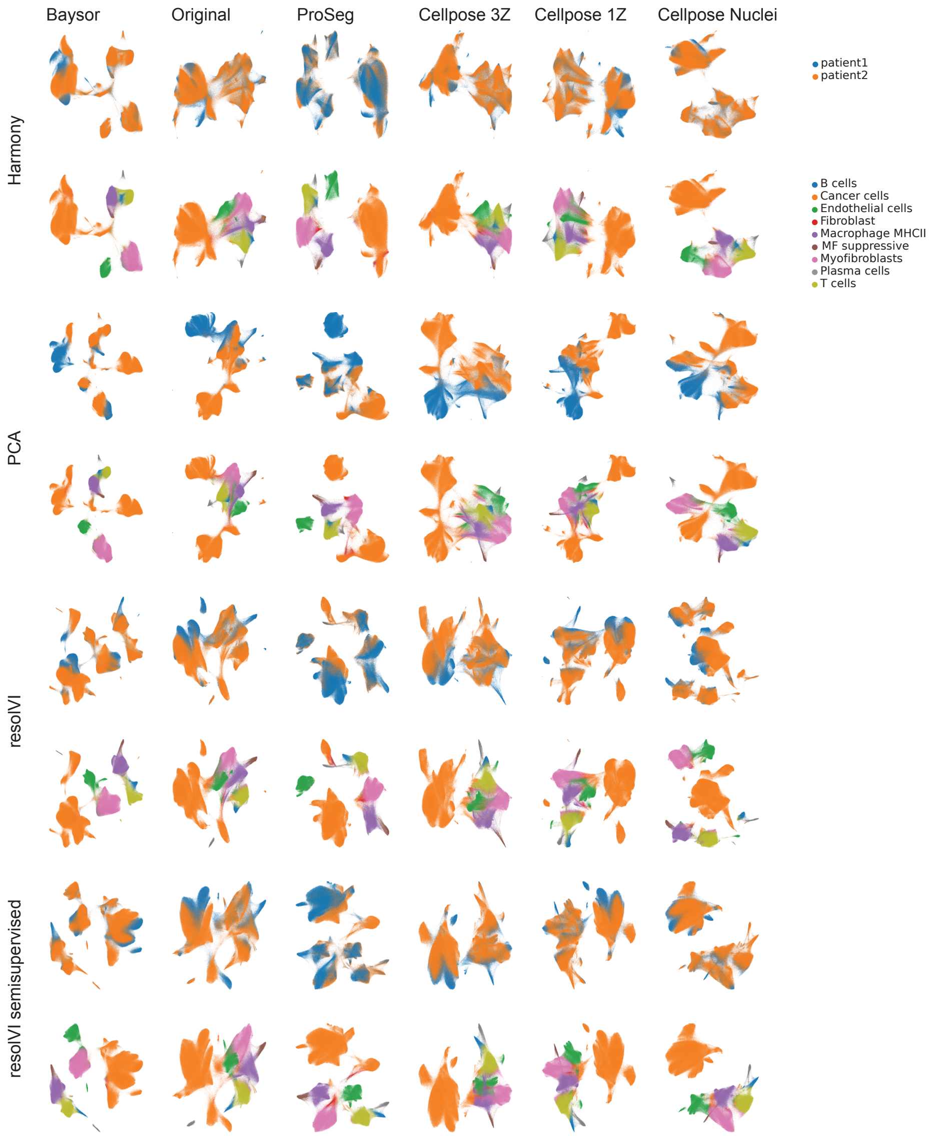


**Supplementary Figure 9: Comparison of UMAP plots for different segmentation algorithms.** We compute UMAP using gene expression computed with different cell segmentation algorithms and four different methods for integration (harmony, PCA, resolVI, and supervised resolVI). For each combination we show two plots one colored by patient ID and one by cell-type. It becomes evident that Baysor and ProSeg UMAP clusters cells by cell-type, while all 3 Cellpose algorithms show much less preservation of cell types.

**Baysor**

| Method | Bio conservation | | | | | Batch correction | | | | Aggregate score | | |
|---|---|---|---|---|---|---|---|---|---|---|---|---|
| | Isolated labels | KMeans NMI | KMeans ARI | Silhouette label | cLISI | Silhouette batch | iLISI | KBET | Graph connectivity | Batch correction | Bio conservation | Total |
| resolVI | 0.64 | 0.81 | 0.90 | 0.62 | 1.00 | 0.90 | 0.11 | 0.43 | 0.91 | 0.59 | 0.79 | 0.71 |
| resolVI supervised | 0.64 | 0.74 | 0.56 | 0.60 | 1.00 | 0.91 | 0.12 | 0.34 | 0.98 | 0.59 | 0.71 | 0.66 |
| PCA | 0.59 | 0.69 | 0.51 | 0.60 | 1.00 | 0.92 | 0.00 | 0.27 | 0.82 | 0.50 | 0.68 | 0.61 |
| Harmony | 0.59 | 0.69 | 0.51 | 0.60 | 1.00 | 0.92 | 0.00 | 0.27 | 0.82 | 0.50 | 0.68 | 0.61 |

**Original**

| Method | Bio conservation | | | | | Batch correction | | | | Aggregate score | | |
|---|---|---|---|---|---|---|---|---|---|---|---|---|
| | Isolated labels | KMeans NMI | KMeans ARI | Silhouette label | cLISI | Silhouette batch | iLISI | KBET | Graph connectivity | Batch correction | Bio conservation | Total |
| resolVI | 0.62 | 0.68 | 0.50 | 0.59 | 1.00 | 0.91 | 0.09 | 0.27 | 0.98 | 0.56 | 0.68 | 0.63 |
| resolVI supervised | 0.61 | 0.64 | 0.56 | 0.61 | 1.00 | 0.90 | 0.06 | 0.25 | 0.96 | 0.54 | 0.68 | 0.63 |
| PCA | 0.57 | 0.55 | 0.35 | 0.59 | 1.00 | 0.91 | 0.00 | 0.15 | 0.86 | 0.48 | 0.61 | 0.56 |
| Harmony | 0.57 | 0.55 | 0.35 | 0.59 | 1.00 | 0.91 | 0.00 | 0.15 | 0.86 | 0.48 | 0.61 | 0.56 |

**ProSeg**

| Method | Bio conservation | | | | | Batch correction | | | | Aggregate score | | |
|---|---|---|---|---|---|---|---|---|---|---|---|---|
| | Isolated labels | KMeans NMI | KMeans ARI | Silhouette label | cLISI | Silhouette batch | iLISI | KBET | Graph connectivity | Batch correction | Bio conservation | Total |
| resolVI | 0.63 | 0.78 | 0.81 | 0.65 | 1.00 | 0.90 | 0.17 | 0.34 | 0.97 | 0.59 | 0.77 | 0.70 |
| resolVI supervised | 0.64 | 0.73 | 0.51 | 0.62 | 1.00 | 0.91 | 0.15 | 0.32 | 0.99 | 0.59 | 0.70 | 0.66 |
| PCA | 0.59 | 0.62 | 0.40 | 0.61 | 1.00 | 0.91 | 0.00 | 0.22 | 0.91 | 0.51 | 0.64 | 0.59 |
| Harmony | 0.59 | 0.62 | 0.40 | 0.61 | 1.00 | 0.91 | 0.00 | 0.22 | 0.91 | 0.51 | 0.64 | 0.59 |

**Cellpose 3Z**

| Method | Bio conservation | | | | | Batch correction | | | | Aggregate score | | |
|---|---|---|---|---|---|---|---|---|---|---|---|---|
| | Isolated labels | KMeans NMI | KMeans ARI | Silhouette label | cLISI | Silhouette batch | iLISI | KBET | Graph connectivity | Batch correction | Bio conservation | Total |
| resolVI | 0.60 | 0.67 | 0.62 | 0.61 | 1.00 | 0.89 | 0.08 | 0.26 | 0.95 | 0.54 | 0.70 | 0.64 |
| resolVI supervised | 0.61 | 0.62 | 0.34 | 0.58 | 1.00 | 0.90 | 0.09 | 0.26 | 0.98 | 0.56 | 0.63 | 0.60 |
| PCA | 0.57 | 0.54 | 0.33 | 0.58 | 1.00 | 0.90 | 0.00 | 0.14 | 0.86 | 0.48 | 0.60 | 0.55 |
| Harmony | 0.57 | 0.54 | 0.33 | 0.58 | 1.00 | 0.90 | 0.00 | 0.14 | 0.86 | 0.48 | 0.60 | 0.55 |

**Cellpose 1Z**

| Method | Bio conservation | | | | | Batch correction | | | | Aggregate score | | |
|---|---|---|---|---|---|---|---|---|---|---|---|---|
| | Isolated labels | KMeans NMI | KMeans ARI | Silhouette label | cLISI | Silhouette batch | iLISI | KBET | Graph connectivity | Batch correction | Bio conservation | Total |
| resolVI | 0.61 | 0.73 | 0.77 | 0.62 | 1.00 | 0.90 | 0.09 | 0.25 | 0.95 | 0.55 | 0.75 | 0.67 |
| resolVI supervised | 0.61 | 0.64 | 0.37 | 0.58 | 1.00 | 0.90 | 0.09 | 0.26 | 0.98 | 0.56 | 0.64 | 0.61 |
| PCA | 0.57 | 0.55 | 0.38 | 0.59 | 1.00 | 0.90 | 0.00 | 0.14 | 0.85 | 0.47 | 0.62 | 0.56 |
| Harmony | 0.57 | 0.55 | 0.38 | 0.59 | 1.00 | 0.90 | 0.00 | 0.14 | 0.85 | 0.47 | 0.62 | 0.56 |

**Cellpose nuclei**

| Method | Bio conservation | | | | | Batch correction | | | | Aggregate score | | |
|---|---|---|---|---|---|---|---|---|---|---|---|---|
| | Isolated labels | KMeans NMI | KMeans ARI | Silhouette label | cLISI | Silhouette batch | iLISI | KBET | Graph connectivity | Batch correction | Bio conservation | Total |
| resolVI | 0.64 | 0.71 | 0.58 | 0.67 | 1.00 | 0.89 | 0.12 | 0.32 | 0.94 | 0.57 | 0.72 | 0.66 |
| resolVI supervised | 0.62 | 0.68 | 0.42 | 0.60 | 1.00 | 0.91 | 0.16 | 0.30 | 0.98 | 0.59 | 0.66 | 0.63 |
| PCA | 0.58 | 0.64 | 0.42 | 0.60 | 1.00 | 0.92 | 0.00 | 0.19 | 0.85 | 0.49 | 0.65 | 0.59 |
| Harmony | 0.58 | 0.64 | 0.42 | 0.60 | 1.00 | 0.92 | 0.00 | 0.19 | 0.85 | 0.49 | 0.65 | 0.59 |

**Supplementary Figure 10: Comparison of scib-metrics for different segmentation algorithms.** We computed scib-metrics for the different segmentations and compare the results here. We find that Baysor and ProSeg perform best in bio conservation, while the original segmentation performs worst in this metric. All segmentations methods performs similarlyin batch correction with a slightly higher score for Baysor and ProSeg. Across all segmentations resolVI and resolVI supervised perform better than PCA and Harmony in both metrics. Highlighting that these methods are superior. ResolVI outperforms the supervised method for most segmentations. Both methods are similar in cLISI and isolated labels, while resolVI mainly outperforms the supervised model in KMeans NMI and ARI. This is likely to higher intra-celltype variation in resolVI supervised for tumor cells, which leads to lower scores in these clustering-based metrics.

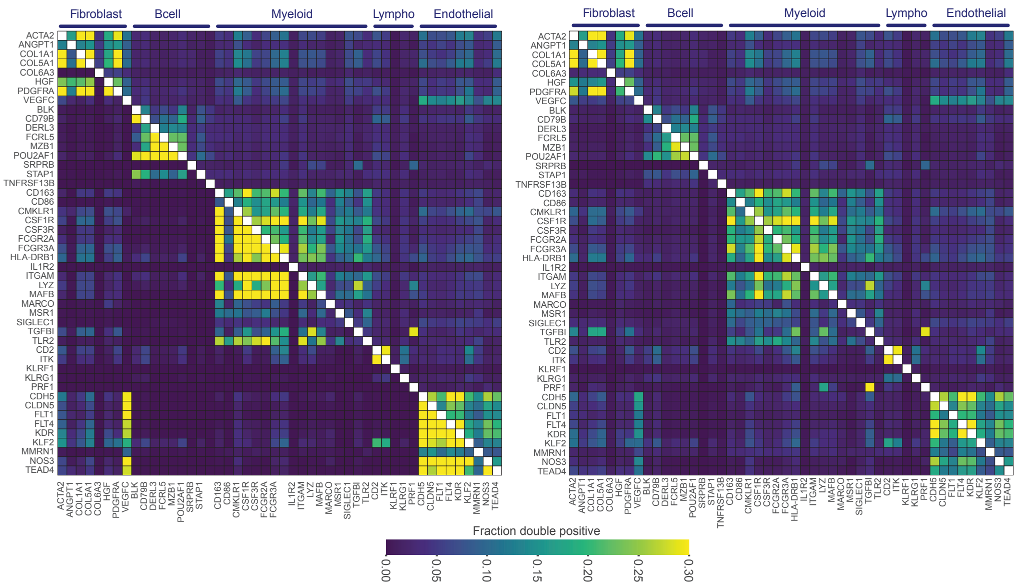


**Supplementary Figure 11: Comparison of fraction double positive for ProSeg segmentation.** Similarly to Figure 3, we compute the fraction of double positive before and after resolVI correction (left) and ProSeg estimation (right). ProSeg can model molecule diffusion and computes a likelihood that a molecule is expressed in a specific cell. We find a clear improvement in fraction of double positive cells for mutually exclusive expressed genes using resolVI correction. In contrast, ProSeg estimated counts provide little improvement over the naively assigned counts. Using ProSeg estimated expression is insufficient in correcting biases in this dataset.

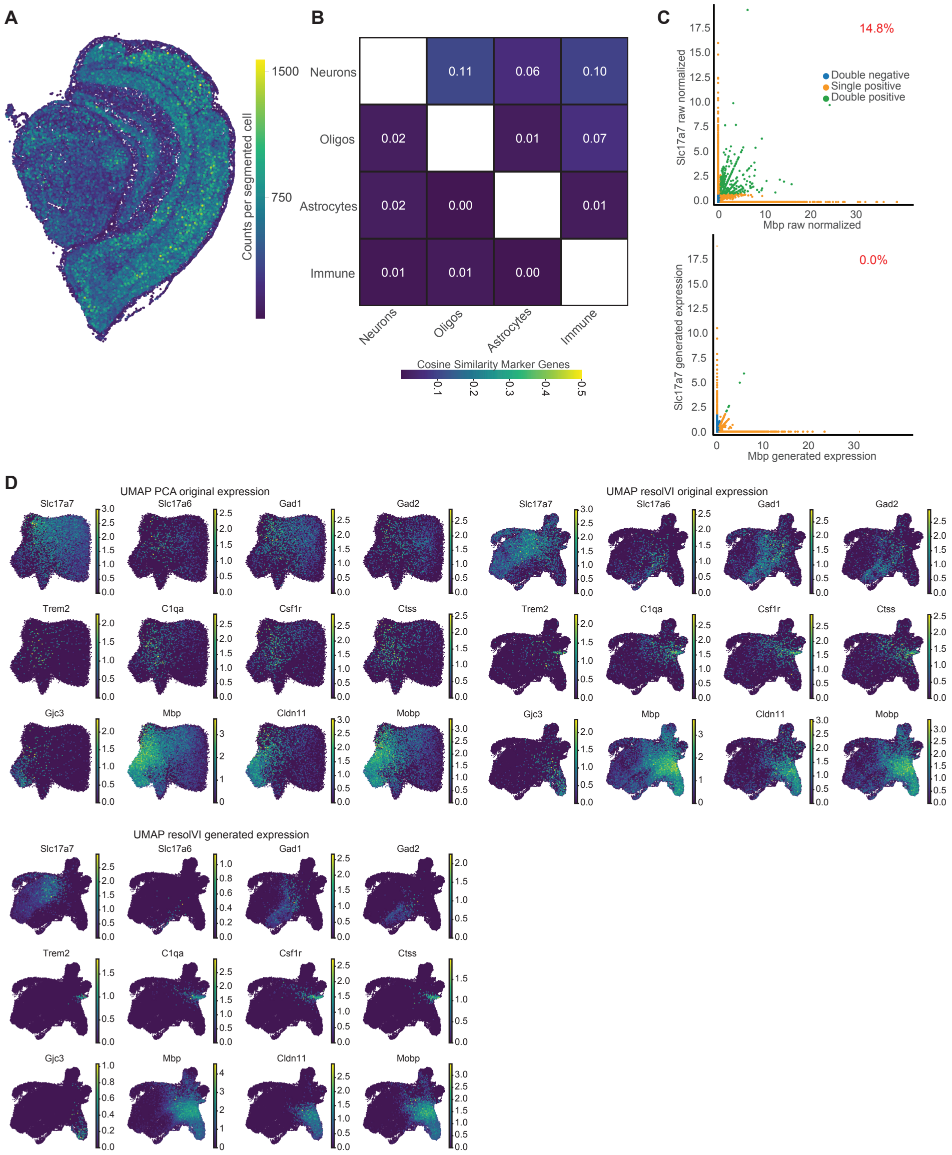


**Supplementary Figure 12: ResolVI improves downstream analysis of Stereo-SEQ data.** We use a demonstration dataset from Stereo-SEQ highlighting a coronal slice of a mouse brain. Cell segmentation was provided by the company. **A** Molecule counts per cell displayed in space. Stereo-SEQ data is much sparser compared to image-based technologies with a limited number of cells with more than 500 counts. **B** Cosine similarity between mutually exclusively expressed marker genes for different cell-types. Counts were library-size normalized and summed for different cell-type markers. Cosine similarity was computed between these summed counts. Above the diagonal we display raw expression values and below the diagonal resolVI generated expression. **C** Quantification of double-positive cells before and after application of resolVI. Displayed are *Slc17a7* (neuron marker) and *Mbp* (oligodendrocyte marker). Percent double positives of *Mbp* single positive cells is displayed. **D** Top left UMAP embedding computed on PCA embedding compared to computed on resolVI latent space. We highlight four markers of neurons (first row), four markers of microglia (second row) and four markers of oligodendrocyte (third row). Top right UMAPs computed on resolVI latent space, we display the raw expression. Bottom right UMAPs computed on resolVI latent space, we display the generated expression.

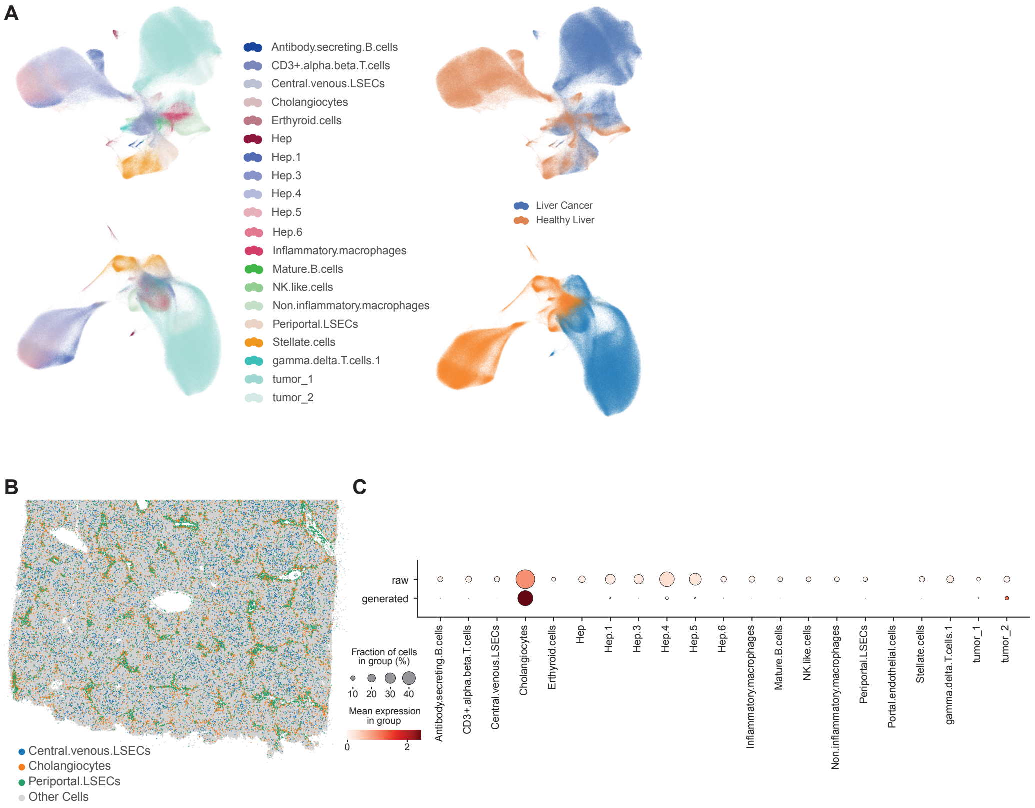


**Supplementary Figure 13: Cholangiocyte distribution and expression in healthy liver. A** UMAP plots in the latent space of resolVI (top) and PCA space computed on original count data. Cell-types are better preserved in the latent space of supervised resolVI. **B** Spatial distribution of cholangiocytes and central venous liver sinusoidal endothelial cells (LSEC). Cholangiocytes show a periportal pattern. **C** Expression of cholangiocyte genes (*SOX4*, *KRT13* and *EPCAM*) across different cell-types is shown. We summed the normalized expression of these genes and display the fraction of cells of a respective type expressing these genes. This signal is specific to cholangiocytes after applying resolVI, while in original counts also other cell-types are positive for these genes.

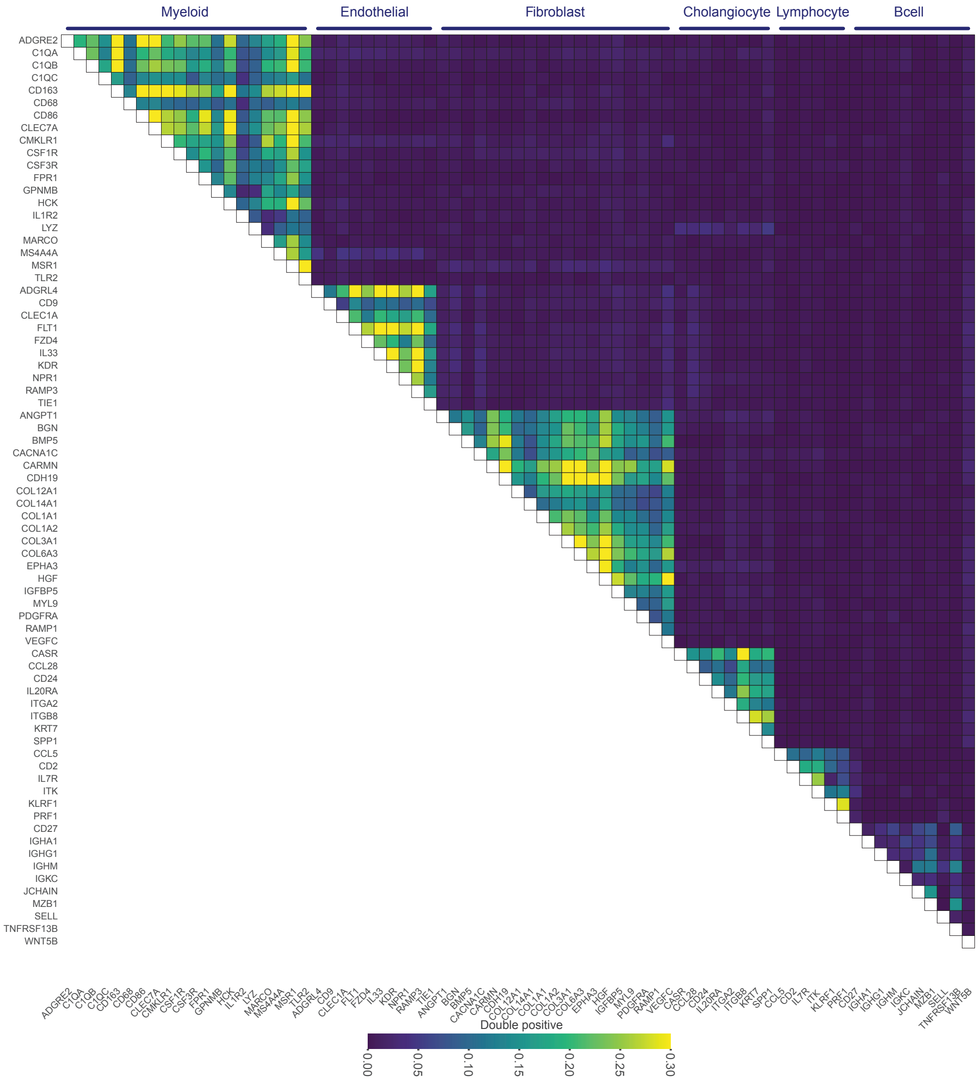


**Supplementary Figure 14: Estimated ratio of double positive expression in single-nucleus reference dataset.** We subset the single-nucleus reference dataset to all genes in the Nanostring panel and compute a Poisson mixture model on this data. Marker genes were selected to be expressed in more than 5% of cells of the respective cell-type and less than 1% in all other cell-types. Displayed is the estimated double positive rate within the original count data of the single-cell nucleus data, while the cell-types are highlighted as the header. We find co-expression of genes within one cell-type, while genes were not explicitly selected to be co-expressed.

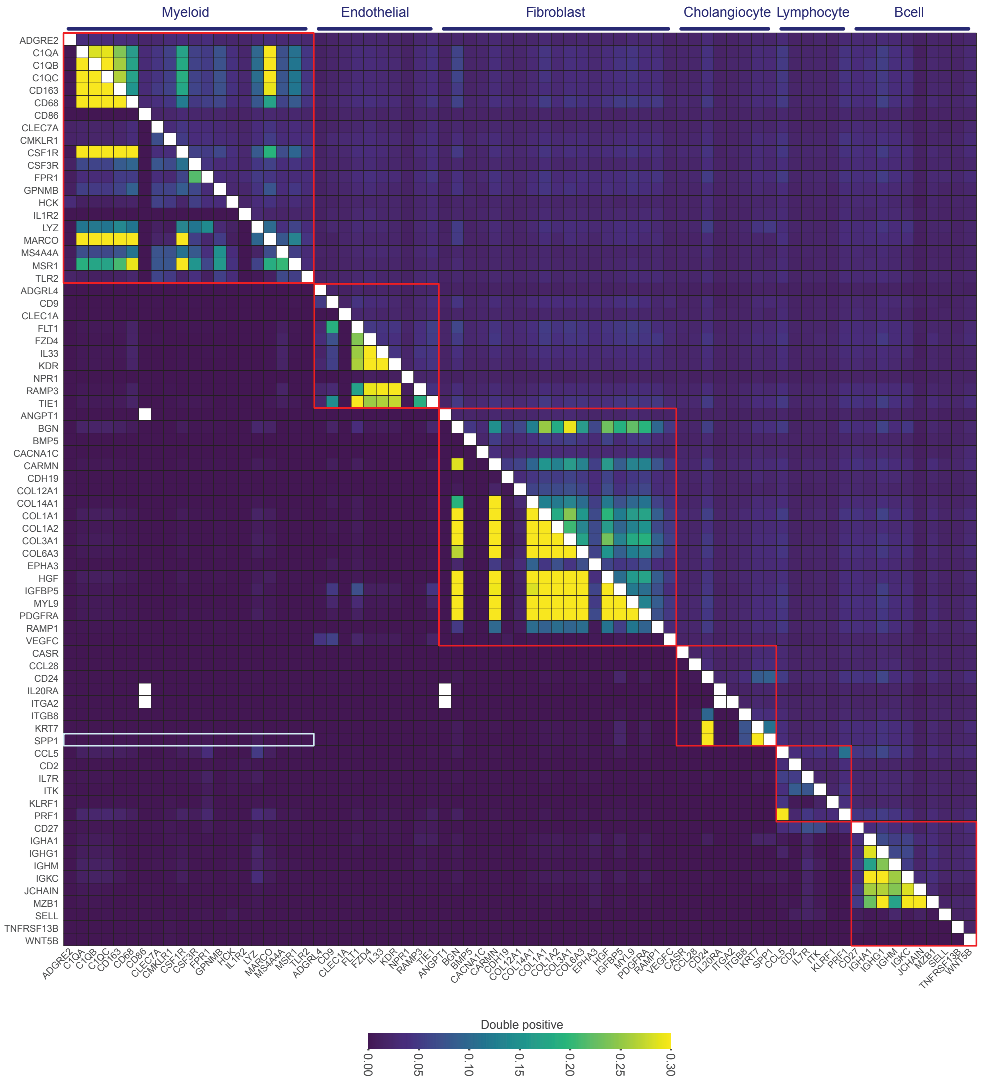


**Supplementary Figure 15: Estimated ratio of double positive expression in healthy liver.** We display the estimated fraction of cells co-expressing a gene pair using a Poisson mixture model. Above the diagonal we display the fraction computed on original expression while below the diagonal we display the results in resolVI generated expression. The red boxes highlight marker genes for the same cell-type, where co-expression is expected. The turqoise box highlights *SPP1* and myeloid cell markers. While these pairs are positive in liver cancer, they are negative in healthy liver tissue. We find a strong reduction in double positive counts of mutually exclusive expressed genes and an increase in co-expression for co-expressed genes. White off-diagonal fields display gene pairs for which the Poisson mixture model predicts both genes to be not expressed after resolVI correction.

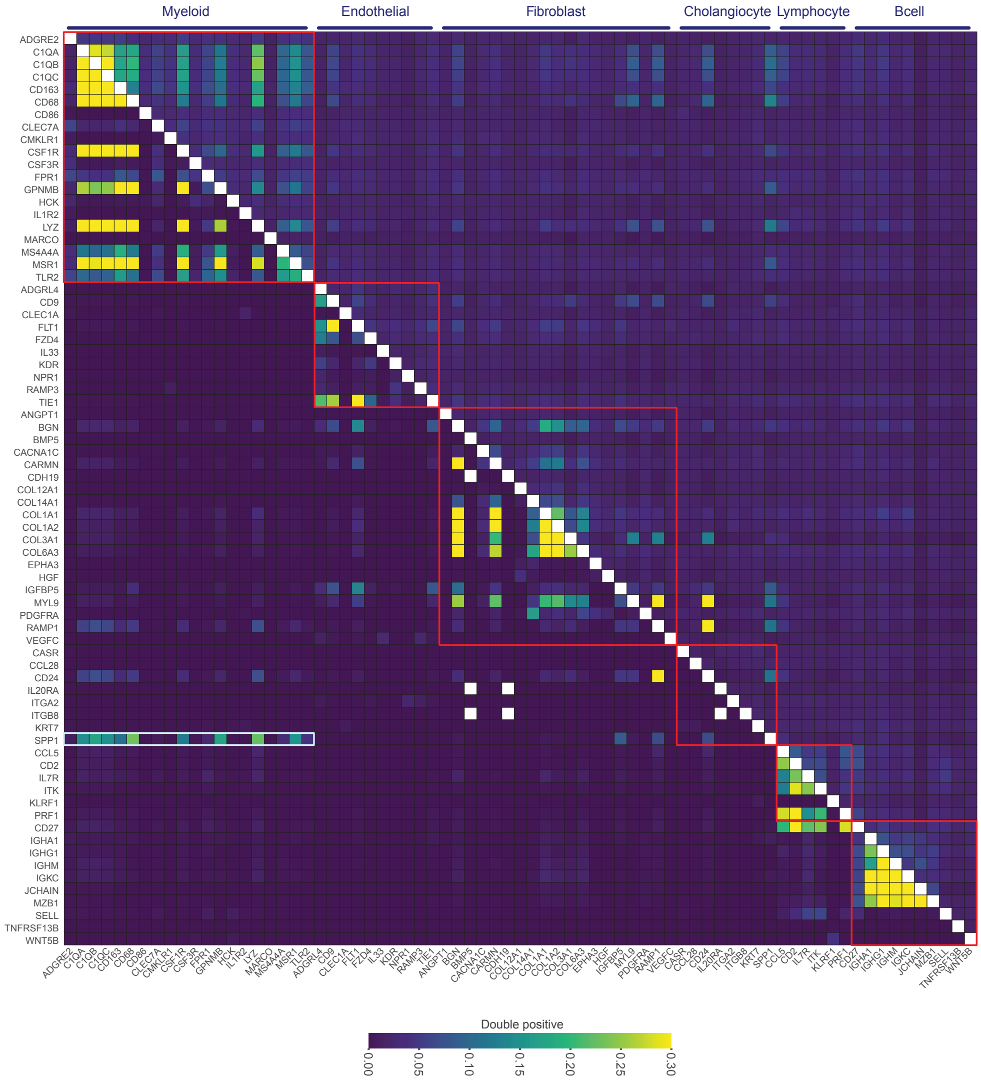


**Supplementary Figure 16: Estimated ratio of double positive expression in liver cancer.** We display the estimated fraction of cells co-expressing a gene pair using a Poisson mixture model. Above the diagonal we display the fraction computed on original expression while below the diagonal we display the results in resolVI generated expression. The red boxes highlight marker genes for the same cell-type, where co-expression is expected. The turqoise box highlights *SPP1* and myeloid cell markers. We find co-expression of these genes in liver cancer. We find a strong reduction in double positive counts of mutually exclusive expressed genes and an increase in co-expression of marker genes of the same cell-type. White fields display gene pairs for which the Poisson mixture model predicts both genes to be not expressed after resolVI correction.

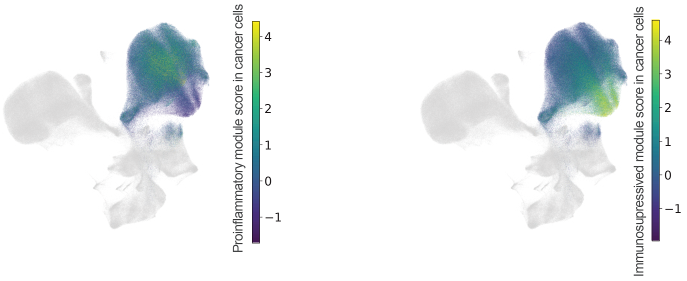


**Supplementary Figure 17: Hotspot modules highlight distinct cancer cells.** Displayed is a UMAP of all cells. We computed the hotspot modules on all cancer cells (all other cells are displayed in grey). Cancer cells are displayed according to their pro-inflammatory module score (left) and anti-inflammatory module score (right).

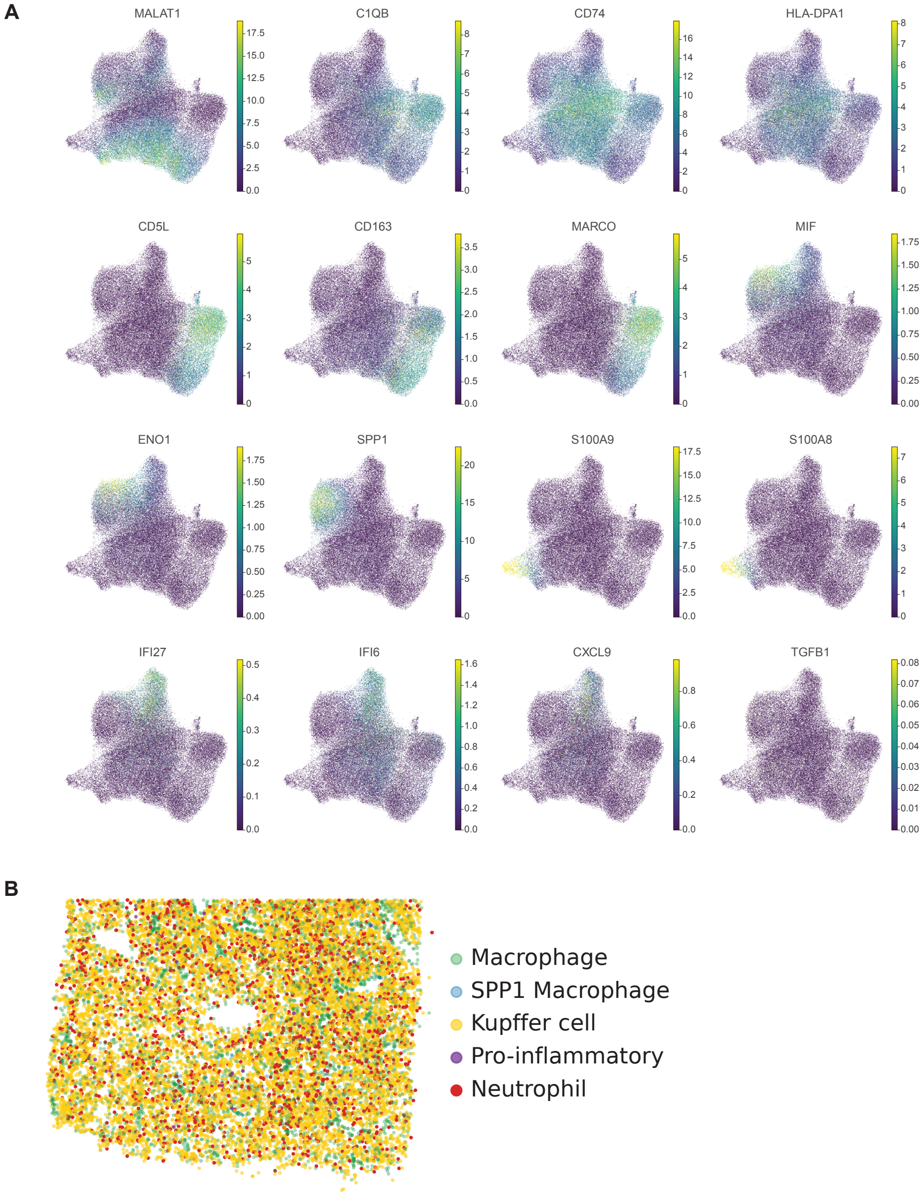


**Supplementary Figure 18: Extended analysis of macrophage cell-types. A** Highlighted are marker genes of distinct macrophage subset overlaid on UMAP plots. We display the resolVI generated counts after count normalization. A clear seperation between the distinct cell-types is visible. Displayed is the generated expression after count-normalization. **B** Spatial distribution of macrophage sub-cell types in the healthy liver highlights neutrophils and Kupffer cells.

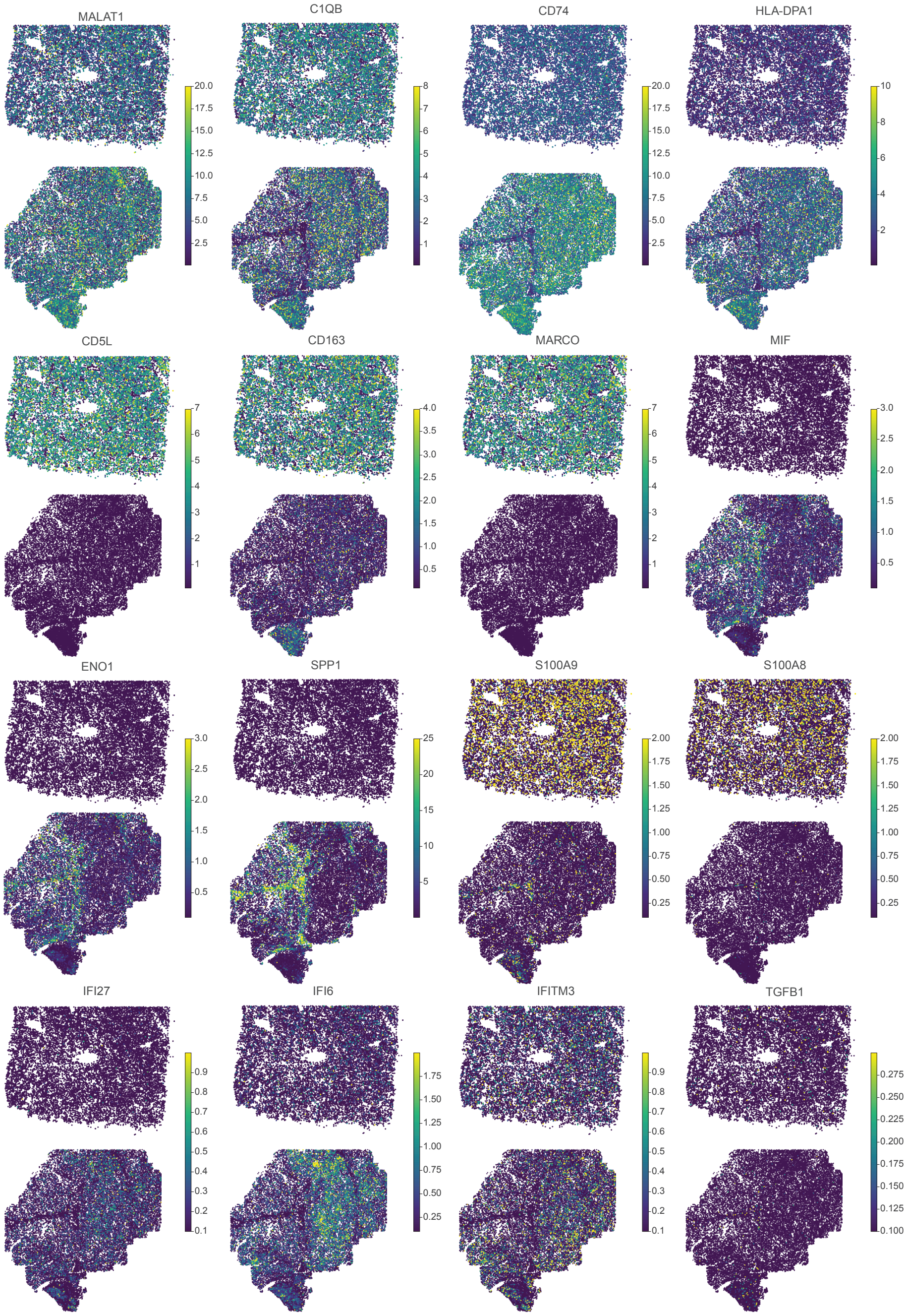


**Supplementary Figure 19: Spatial analysis of macrophage cell-types.** We display marker genes of macrophage subsets in space. Displayed is only the resolVI generated expression within myeloid cells and after count normalization. The upper image each is healthy liver while the lower image is liver cancer. Kupffer cell genes are only visible in the healthy liver. Anti-inflammatory macrophages are only visible in the region annotated as anti-inflammatory tumor cells. Neutrophil genes are visible in the healthy liver and in liver cancer only in the anti-inflammatory region. Interferon-related genes are increased in liver cancer in the pro-inflammatory region. *TGFBI* is lowly expressed but is expressed in the center of the anti-inflammatory region.

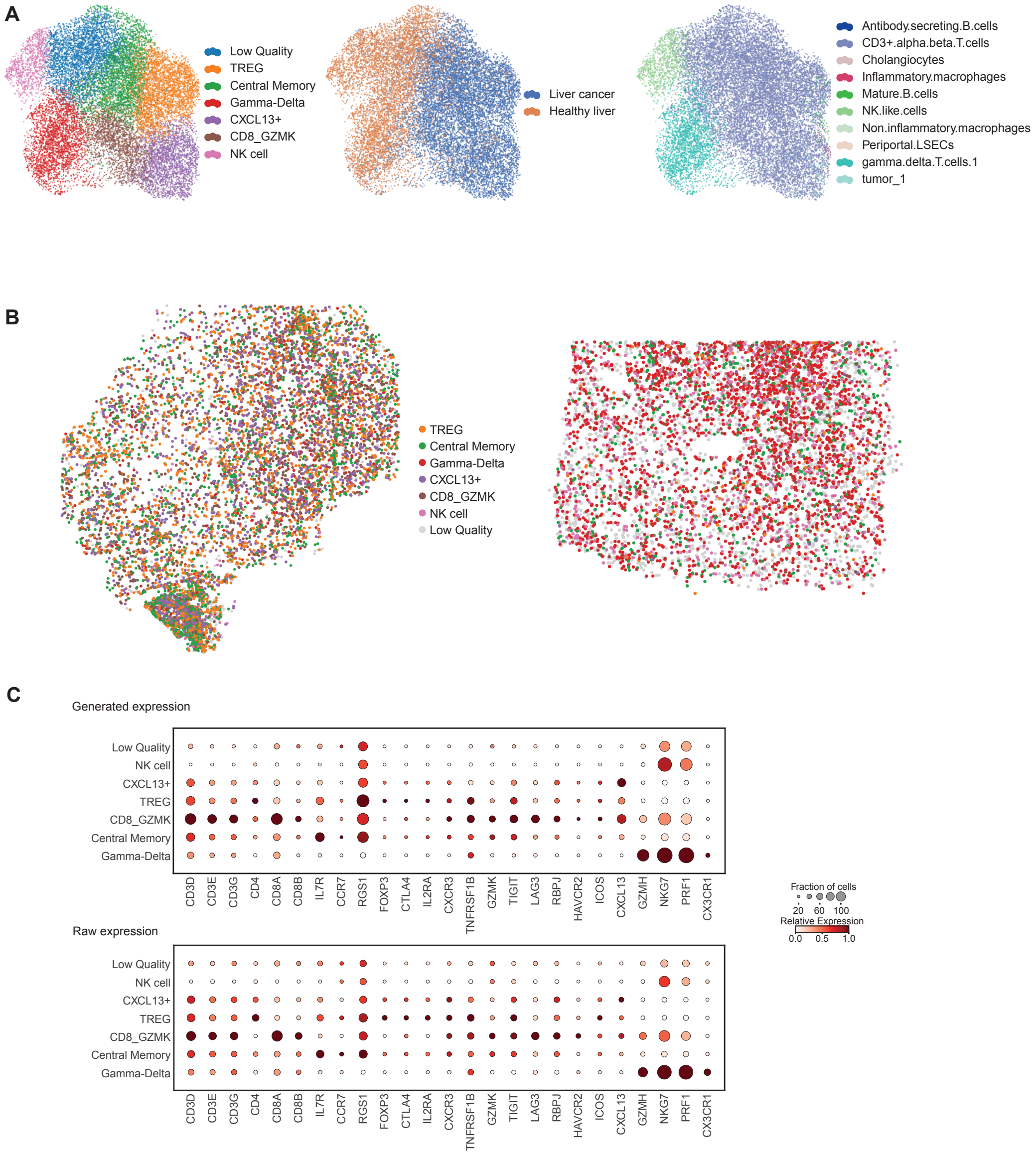


**Supplementary Figure 20: Analysis of T cells in liver cancer (CosMx) data. A** We performed sub-clustering of T cells, while Nanostring annotated NK cells, Gamma-Delta T cells and Alpha-Beta T cells (right). Further investigation after resolVI embedding reveals CXCL13+ CD8 T cells, Central Memory T cells, TREG and CD8 GZMK+ T cells (left). T cell sub-cell types are almost exclusive in either liver cancer or healthy liver. **B** Spatial distribution of the distinct cell-types in liver cancer (left) and healthy liver (right). We find a reduction of T cell in the immuno-suppressive region, while in the lower left a dense aggregate of T cells is visible. **C** We compare manually selected marker gene expression after normalization in raw expression as well as after resolVI correction. Both dotplots show a similar pattern, highlighting that resolVI learns the co-expression pattern of these finely resolved cell-types.

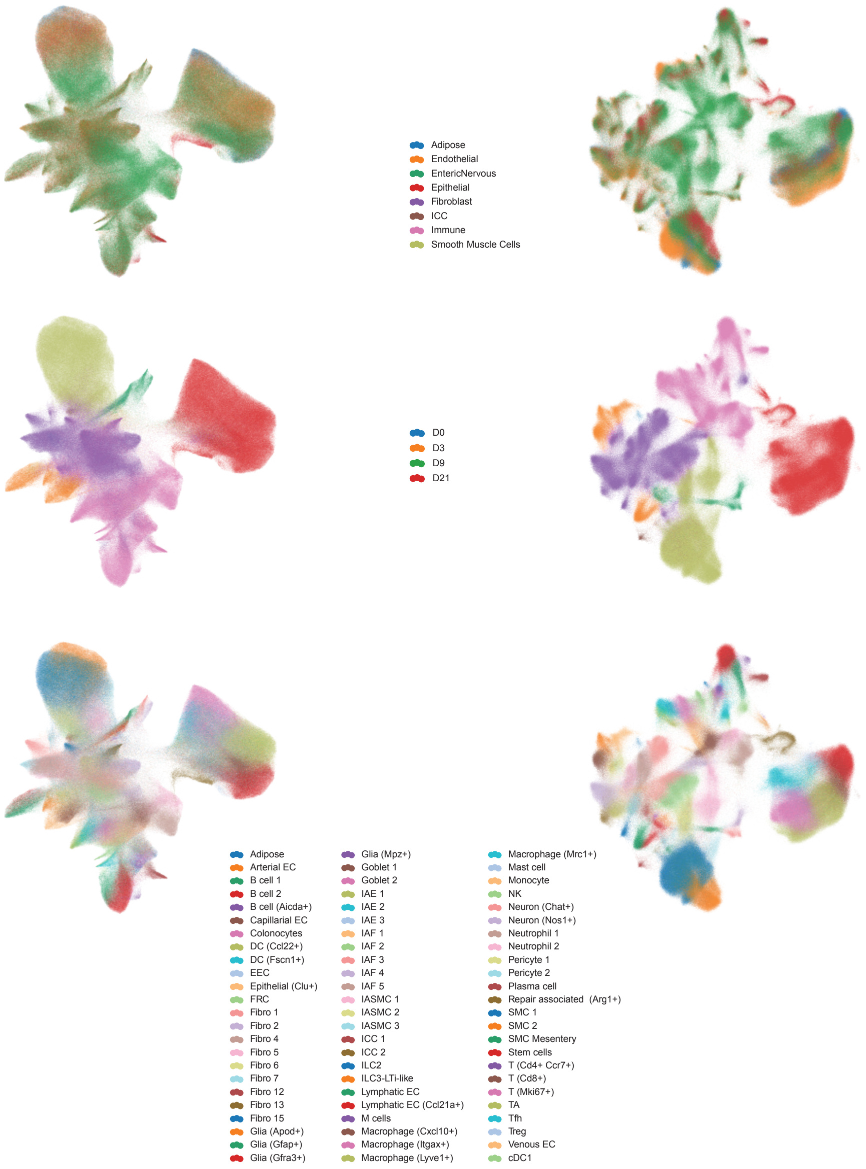


**Supplementary Figure 21: Comparison of PCA on raw expression and resolVI latent space based UMAP embedding of DSS colitis data set.** UMAP of DSS colitis data set colored by time point (top row), coarse cell-types (middle row) and finest cell-type (bottom row). On the right side the embedding was computed after resolVI embedding without cell-type supervision, while the left side shows the original author provided embeddings based on PCA of normalized expression data. Especially for smooth muscle cells and stem cells the resolVI embedding shows more integration.

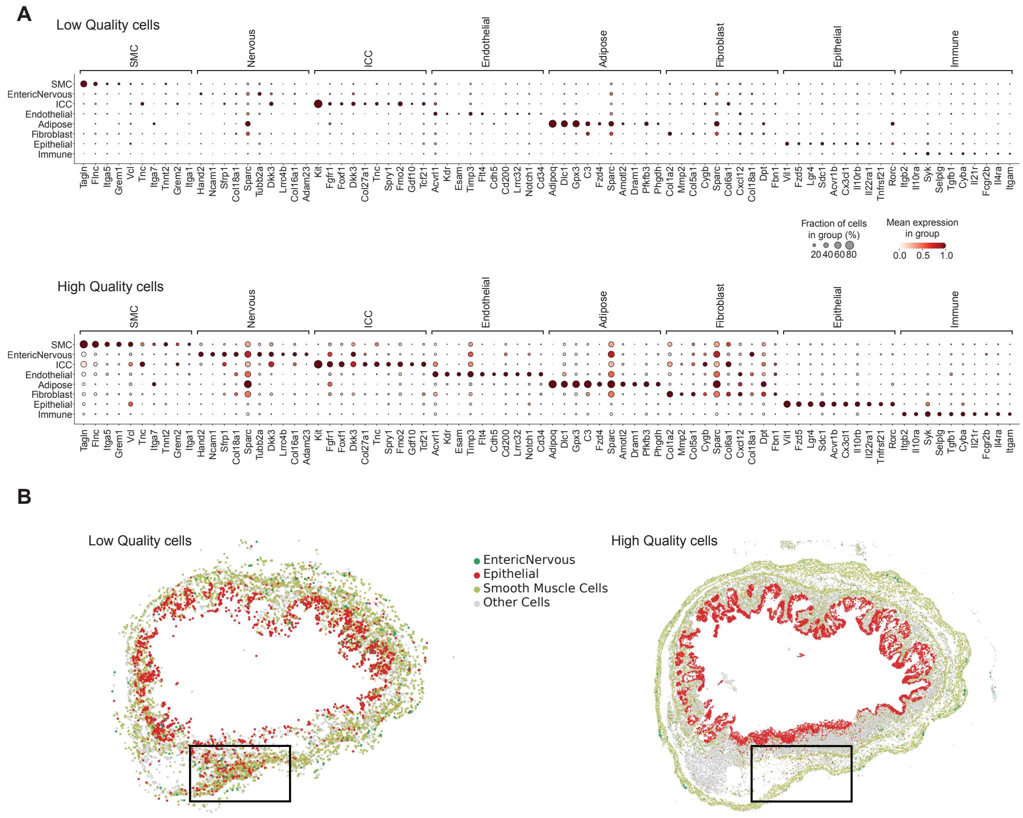


**Supplementary Figure 22: Low quality cells can be detected by expression and location. A** We highlight here cells with estimated true counts by resolVI below 20 (top plot) and all other cells (bottom plot). We performed differential expression using scanpy Wilcoxon test between the coarse cell-types. Dotplot is showed seperately for high and low quality cells. We show the raw count-normalized expression of these cells. In the upper plot for some cell-types no clear marker gene expression is visible like epithelial, endothelial and ervous, whereas in the lower plot a relevant number of cells express the selected marker genes. **B** Spatial plot for a slice at day 9 with many low quality cells. We show separately the location of low (left) and high (right) quality cells. We find in low quality cells many epithelial cells in the muscle layer (black box). This does not fit to the expected anatomical location of these cells (luminal) and for high quality cells we find no cells in this location. A high number of epithelial cells in the low quality cells still highlights regions in the epithelial layer and therefore have the correct location.

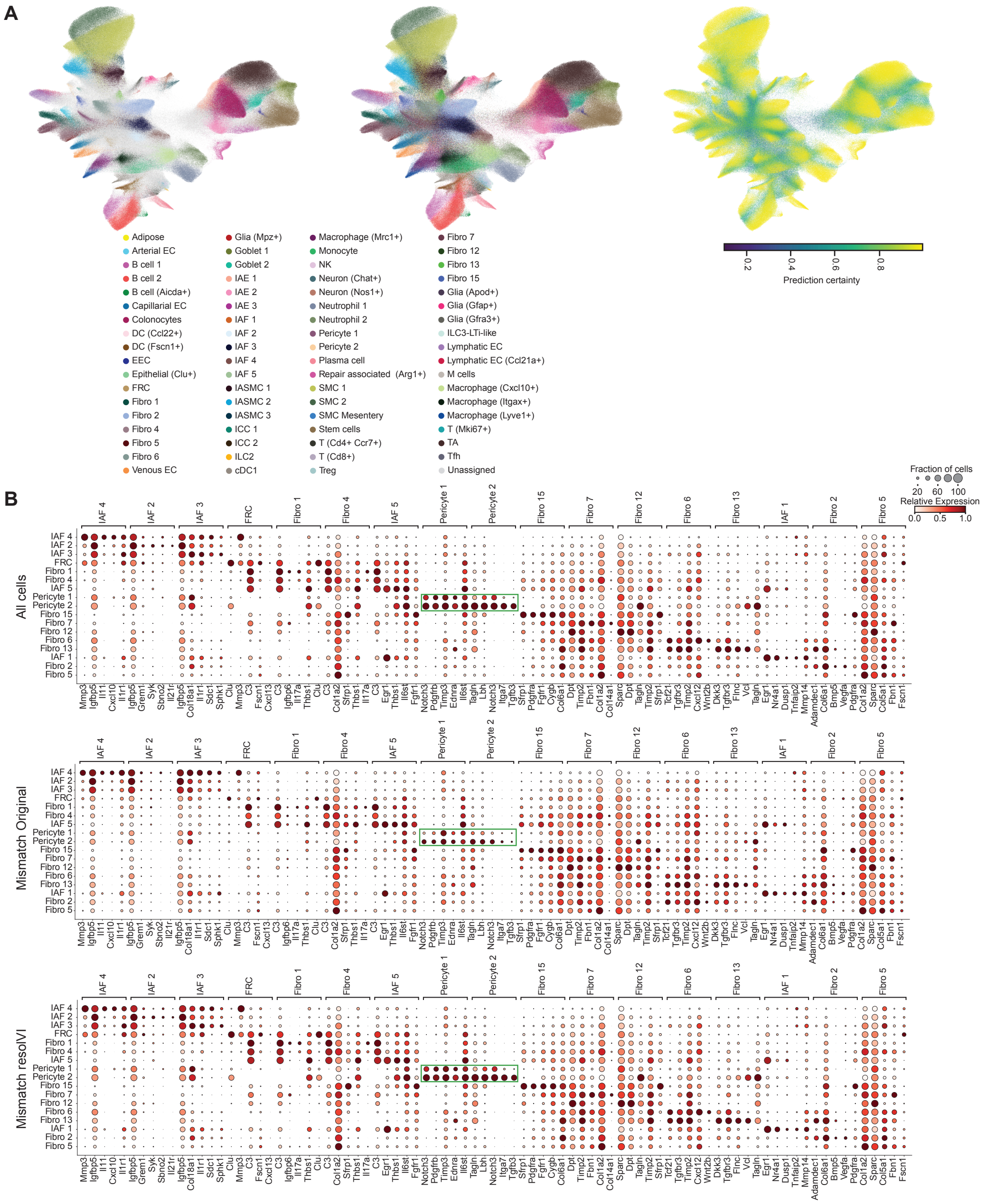


**Supplementary Figure 23: Reassignment of cell-types with resolVI. A** UMAP embedding based on the semi-supervised resolVI model. Fine cell-type annotation after removing labels from cells with low purity in their neighborhood (Methods, right plot cells labeled unassigned are grey), after label transfer using supervised resolVI model (middle) and prediction uncertainty in resolVI (right). **B** Marker genes of all fibroblast cell-types computed on all cells (top), cells with different label after resolVI and author provided (Mismatch) grouped by their original label (middle) and by their resolVI label (lower). Overall, the labels provided by resolVI better agree with the marker genes computed on all cells (highlighted by green box for pericytes).

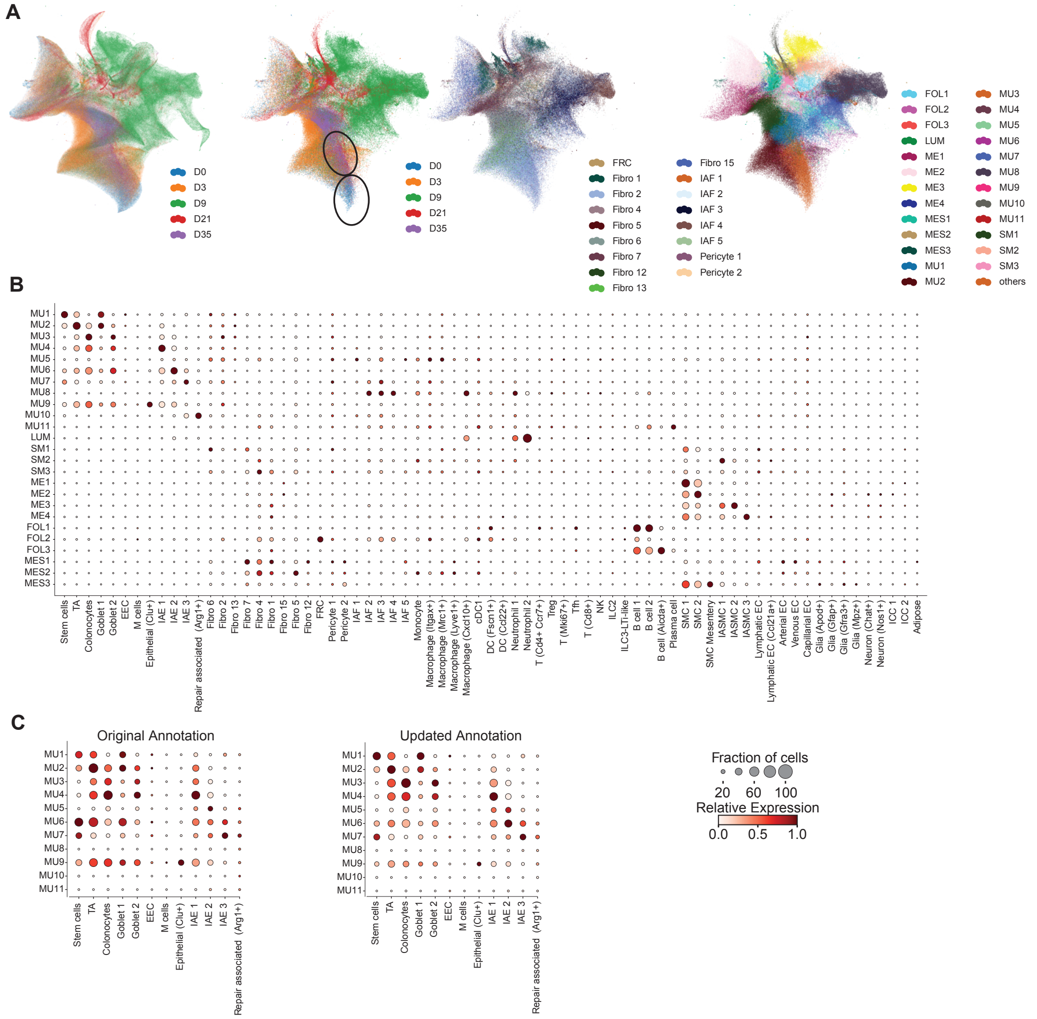


**Supplementary Figure 24: ResolVI improves spatial niche embedddings. A** We computed an embedding using the bag of classifier probabilities (see method). In the left plot we highlight the different time points for all cells, in the other plots we subset the display to all fibroblasts. We display the time points in the second plot, the fine cell-type labels in the third plot and the niche annotation in the fourth plot. In the second plot, we highlight the region enriched for cells from day 35 and cells from day 0. In these regions most fibroblasts are annotated as fibroblast 2. This is why we focus in the main figure on fibroblast 2. We performed $k$-nearest-neighbor classification to propagate the original niche labels to this new embedding. **B** We display the neighborhood composition for each niche annotation and cell-type. **C** For all epithelial regions with a different label after resolVI propagation and the original annotation, we display the cell-type fraction in each region. We find that the updated annotations better reflect the composition of all spatial niches (MU1 is high in colonocytes, MU2 in TA cells and MU3 in stem cells).

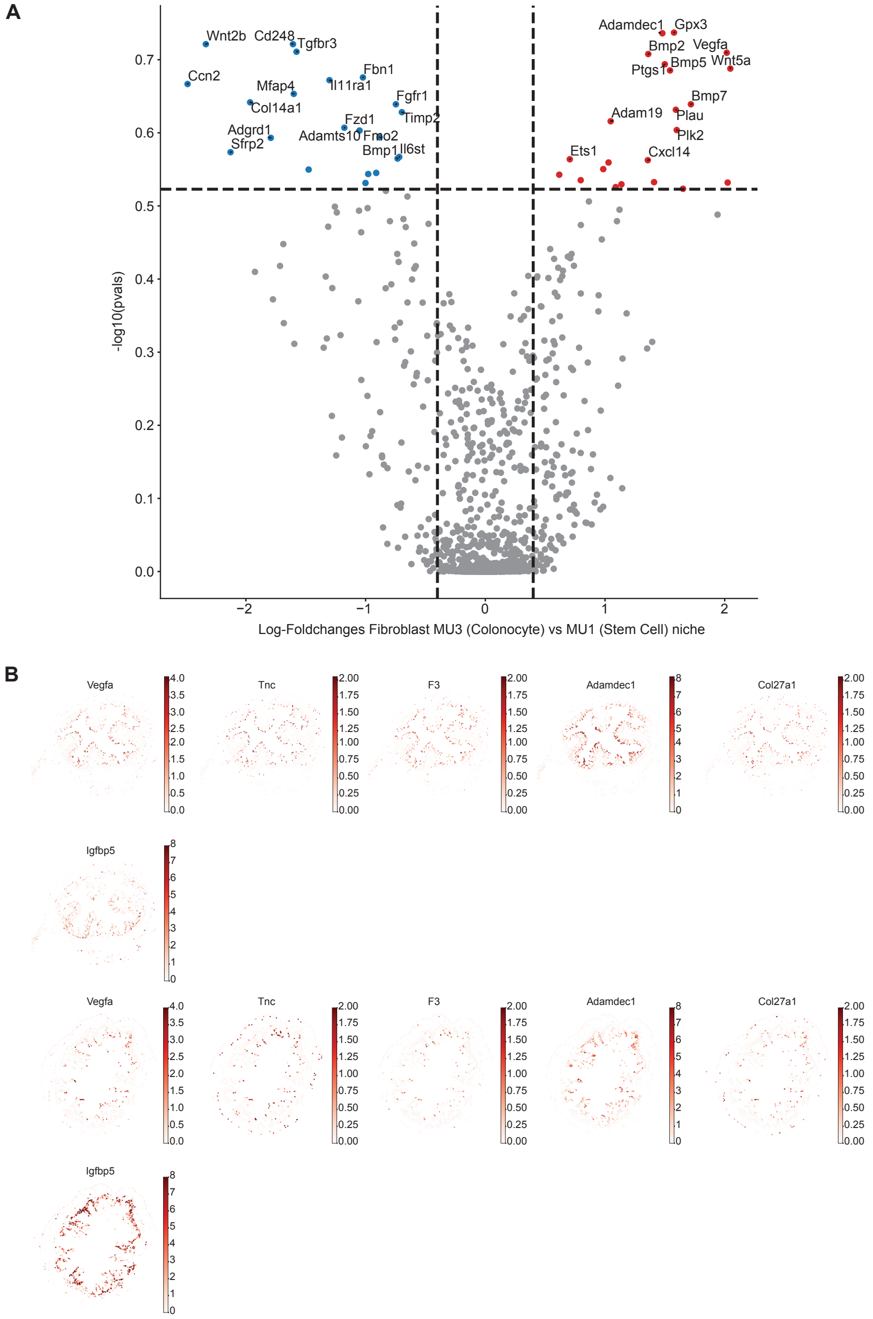


**Supplementary Figure 25: Differential expression between fibroblast 2 in distinct locations. A** We performed differential expression analysis using resolVI between fibroblast 2 cells in the spatial niche MU1 versus MU3. Upregulated genes are higher expressed towards the tip of the crypt while downregulated genes are expressed higher towards the base of the crypt. We find upregulation of Bmp related genes. **B** Spatial display of differentially expressed genes at day 0 (top rows) and day 35 (bottom rows). Indeed, we find higher expression of *Vegfa*, *Tnc*, *F3* and *Col27a1* at day 0, which contains more fibroblasts in the tip of the crypt. And higher expression of *Igfbp5* and *Tgfbr3* at the base of crypt at day 35.

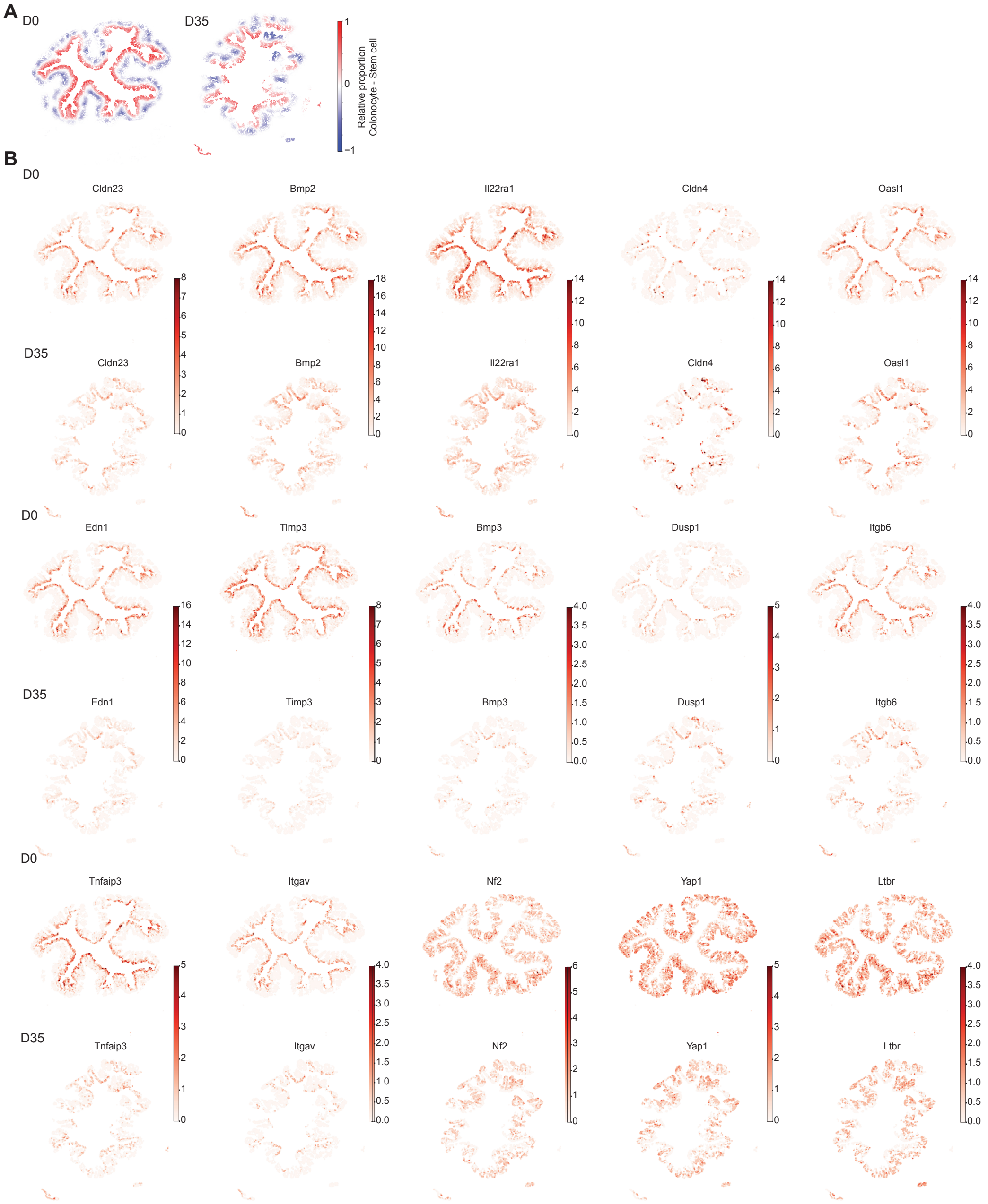


**Supplementary Figure 26: Differential expression between epithelial cells in distinct neighborhoods. A** We display the difference in relative proportion of stem cells and colonocytes at day 0 and day 35 (same display as in Figure 5 but displayed for the whole slice). **B** Apart from the last three plots, genes are upregulated in neighborhoods with a high fraction of colonocytes. Indeed, we find these genes higher expressed at day 0 and find them specifically expressed towards the intestinal lumen. Genes higher expressed in neighborhoods with view colonocytes (last 3 plots are genes that are expressed across the intestinal lumen. The expression is similar at day 0 and day 35.
